# Determining molecular archetype composition and expression from bulk tissues with unsupervised deconvolution

**DOI:** 10.1101/2021.07.12.452047

**Authors:** Chiung-Ting Wu, Lulu Chen, David M. Herrington, Minjie Shen, Guoqiang Yu, Robert Clarke, Chunyu Liu, Yue Wang

## Abstract

Complex tissues are composite ecological systems whose components interact with each other to create a unique physiological or pathophysiological state distinct from that found in other tissue microenvironments. To explore this ground yet dynamic state, molecular profiling of bulk tissues and mathematical deconvolution can be jointly used to characterize heterogeneity as an aggregate of molecularly distinct tissue or cell subtypes. We first introduce an efficient and fully unsupervised deconvolution method, namely the Convex Analysis of Mixtures – CAM3.0, that may aid biologists to confirm existing or generate novel scientific hypotheses about complex tissues in many biomedical contexts. We then evaluate the CAM3.0 functional pipelines using both simulations and benchmark data. We also report diverse case studies on bulk tissues with unknown number, proportion and expression patterns of the molecular archetypes. Importantly, these preliminary results support the concept that expression patterns of molecular archetypes often reflect the interactive not individual contributions of many known or novel cell types, and unsupervised deconvolution would be more powerful in uncovering novel multicellular or subcellular archetypes.

## 1 Introduction

The functions of complex tissues are orchestrated by a productive interplay between many specialized tissue, cell, and task subtypes (Lake, et al., 2016). Characterizing the presence and dynamics of these components is important to understanding many physiological or pathophysiological processes. For example, shifts in the relative composition of neuron or glia cell subtypes is central to the developmental processes of the human brain (Colantuoni, et al., 2011; Kang, et al., 2011). Likewise, understanding the genuine changes of subtype-specific molecular expressions is of direct etiological interest for many diseases (Herrington, et al., 2018). Biologists have amassed a large body of information about complex tissues and have some powerful insights into how they remodel under different conditions (Dadgar, et al., 2014). However, most of our knowledge relies on mixed readouts with many unknown confounders. Experimental solutions to mitigate tissue heterogeneity are to isolate individual cells or to microdissect tissue subtypes before molecular profiling. While promising, physical separation is clearly not the most reliable and cost-effective method and is inapplicable to previously-assayed samples (Avila Cobos, et al., 2018). Would it not be better if we could frame a tissue ecosystem in precise mathematical models and use computational unsupervised deconvolution to analyse, or re-analyse, the wealth of publicly available multi-omics bulk data?

The primary objective of mathematical deconvolution is to computationally detect subtype-specific markers, determine the number of constituent subtypes, calculate subtype proportions in individual samples, and estimate subtype-specific expression profiles (Avila Cobos, et al., 2018; Chen, et al., 2020; Wang, et al., 2016). Complex tissues – where formation and remodeling require productive interactions between multiscale subtypes – are intrinsically dynamic. Thus supervised methods are poorly suited to finding molecularly distinctive subtypes that are subtle, condition-specific (their distinctive signatures change when the cells are present in different microenvironments), or previously unknown (Houseman, et al., 2016; Kuhn, et al., 2011; Newman, et al., 2015). Unsupervised deconvolution methods, which are based on regularized matrix factorization, are an appropriate tool to characterizing the molecular landscape of complex tissues. Supported by advanced machine learning algorithms and proven theorems, unsupervised deconvolution methods can decompose the mixed molecular signals into many latent variable; these subtypes are biological interpretable and functionally enriched (Hart, et al., 2015; Herrington, et al., 2018; Moffitt, et al., 2015; Wang, et al., 2016). However, the question remains as to whether it is possible to develop fully unsupervised deconvolution methods that can concisely define molecular latent variable ecosystem of complex tissues. In this article, we first briefly introduce a fully unsupervised deconvolution method, namely Convex Analysis of Mixtures (CAM1.0-3.0) (Chen, et al., 2020; Wang, et al., 2016). We then report diverse case studies on multi-omics data obtained from bulk tissues, where the type, number, proportion and profile of the molecular components are unknown. Last, we provide an outlook into future methodological development directions.

Our deconvolution pipeline is built on the strong parallelism between linear latent variable models and the theory of convex sets (**Fig. 1**) (Chan, et al., 2008; Chen, et al., 2011; Wang, et al., 2016). Tissue samples to be analyzed by CAM contain an unknown number and varying proportions of molecularly distinctive subtypes (including novel) present across various scales (tissue, cell, or archetype) (**Fig. 1a**). Molecular expression in a specific subtype is modeled as being linearly proportional to the abundance of that subtype. Applying the newly-proven mathematical theorems (Wang, et al., 2016), we showed that the simplex of mixed expression patterns in bulk tissues, in the scatter space of the mixtures (for example, gene expression scatter space), is a rotated and compressed version of the simplex of subtype expressions (**Fig. 1d**). According to the theory of convex sets (Chan, et al., 2008), every molecular feature within the scatter simplex can be uniquely determined by the linear combination of the vertices. Thus, the number of the vertices corresponds to the number of molecularly distinctive subtypes present in the bulk samples and the molecular features residing at the vertices are the molecular markers defining such subtypes (Wang, et al., 2016).

**Figure 1.**
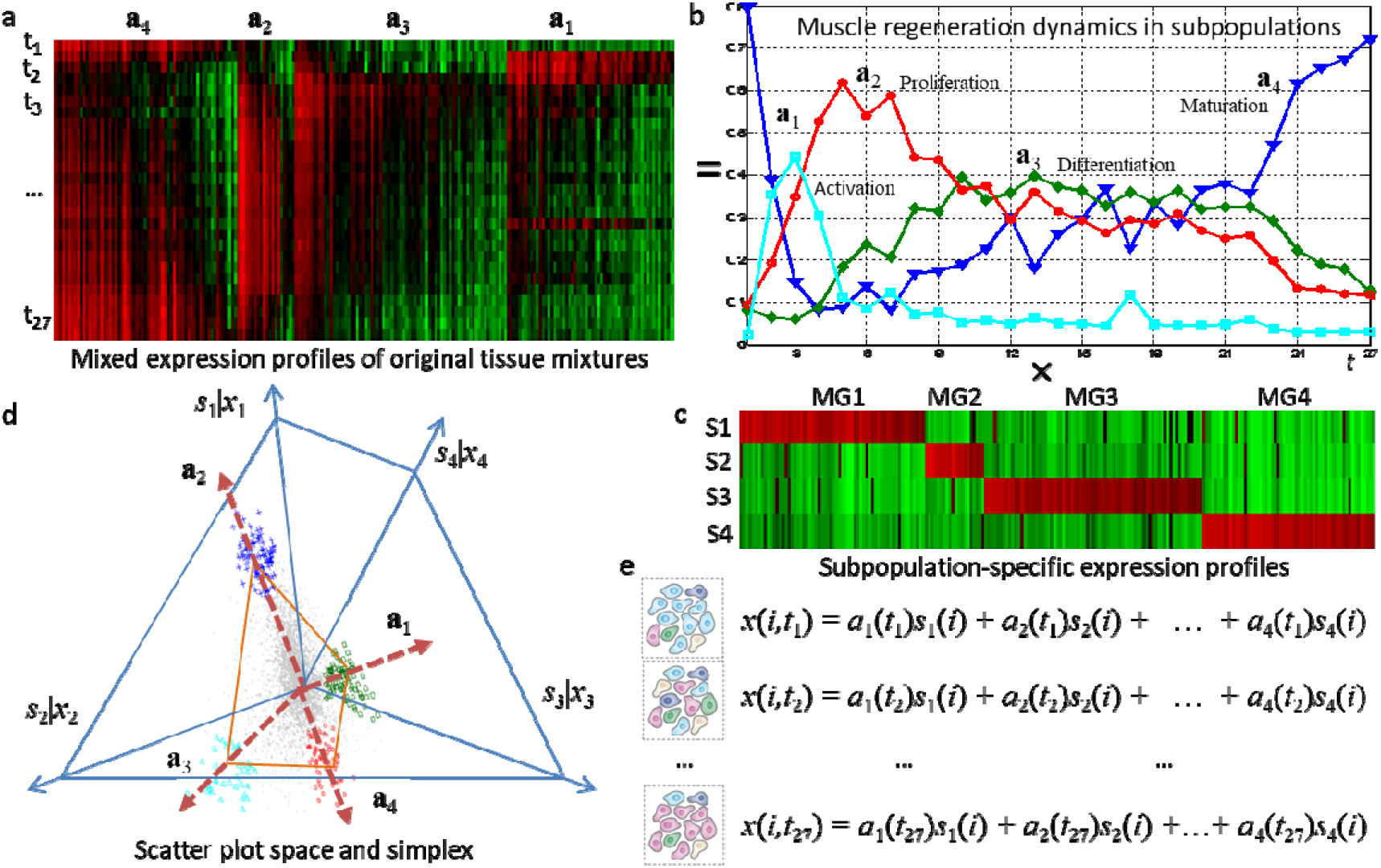
CAM framework for unsupervised *in silico* deconvolution of tissue heterogeneity that uncovers *de novo* subpopulations and repopulation dynamics. In CAM model, *s_k_*(*i*) is the expression level of gene *i* in subpopulation *k*, *x_j_*(*i*) is the expression level of gene *i* in heterogeneous sample *j*, and *a_jk_* is the proportion of subpopulation *k* in heterogeneous sample *j*. In our mathematical modeling of transcriptional heterogeneity, mRNA expression levels from a mixture of multiple subpopulations are modeled as the weighted average of the expression levels from a set of distinct subpopulations, where the weights describe the proportions of each distinct subpopulation in the overall population. (**a**) Heatmap of mixed gene expression time-course from heterogeneous muscle tissues sampled across 27 time points during normal muscle regeneration, plotted over 214 CAM-detected subpopulation-specific marker genes whose expressions are exclusively enriched in a particular subpopulation. (**b**) Repopulation dynamics of the four distinct subpopulations/phases (Activation, Proliferation, Differentiation, and Maturation) in terms of relative individual proportions estimated by CAM. (**c**) Heatmap of marker gene expression levels in the four distinct pure subpopulations/phases estimated by CAM. (**d**) Geometry of the mixing operation in scatter space that produces a compressed and rotated scatter simplex of original mixed gene expressions whose vertices host subpopulation-specific marker genes and correspond to mixing proportion vectors. (**e**) Mathematical description on the mixed gene expression readout from heterogeneous mixtures of multiple distinct subpopulations.

Our CAM3.0 based deconvolution works by detecting the vertices of the scatter simplex geometrically, *i.e*., determining the multifaceted simplex that most tightly encloses the globally measured expression mixtures. Subsequently, we identify the molecular markers residing at the vertices, and estimate the proportions and specific expression profiles of constituent subtypes (Wang, et al., 2016). The number of subpopulations present is determined by the newly-derived minimum description length (MDL) criterion. Ideally, a molecularly distinctive subtype would contain molecular signatures (molecular markers) that are exclusively expressed in the cognate cell or tissue subtype of interest while in no others. Importantly, our deconvolution pipeline requires no a priori information on the number, signatures, or compositions of the subtypes present in heterogeneous samples, and does not require the presence of pure subtype samples (Hart, et al., 2015; Schwartz and Shackney, 2010). This advantage is significant in that CAM3.0 can achieve all of its goals using only a small number of heterogeneous samples, and provides a powerful means to distinguish among phenotypically similar subtypes.

While there may be many ways to subdivide a complex tissue ecosystem, there is clearly a need to dissect the staggering complexity of many multiscale molecular landscapes. We have shown the performance and biomedical utility of CAM3.0 based deconvolution using gene expression (Wang, et al., 2016), methylation (Chen, et al., 2020), proteomics (Herrington, et al., 2018; Parker, et al., 2020), and imaging data (Chen, et al., 2011; Fan, et al., 2020). These applications have led to novel findings and hypotheses. A molecular latent variable model of complex tissues, particularly in the presence of disease lesions, is not yet available, but we can provide a roadmap of how a machine learning approach might uncover, in mathematical forms, the molecular events controlling tissue remodeling in many biomedical contexts. The value of this deep characterization is illustrated by the fully unsupervised deconvolution results obtained from spatial-temporal molecular expression data of various complex tissues, and will be measured ultimately by the emerged new insights or hypotheses.

## 2 Results

We have developed the open-source Bioconductor R package that implements and tests the basic functionalities of the CAM1.0/2.0 deconvolution pipeline (Chen, et al., 2021; Chen, et al., 2020), freely available at http://bioconductor.org/packages/debCAM. Experimental validation and comparison on CAM1.0/2.0 method with most relevant peer methods have been previously reported (Chan, et al., 2008; Chen, et al., 2020; Wang, et al., 2010; Wang, et al., 2016; Zhu, et al., 2016). The CAM3.0 software tool is freely available at https://github.com/ChiungTingWu/CAM3

### Muscle regeneration

This case study aims to investigate whether and how the interactions among the molecular subtypes may affect the normal or failed (muscular dystrophies) muscle regeneration. Because muscular dystrophy remodeling is expected to share many features with normal muscle regeneration (**Fig. 1**), we sought to determine the molecular mechanisms underlying observed failure of muscle regeneration in dystrophin-deficient muscle through hypothesis generation using muscle mRNA profiling data (Bakay, et al., 2006; Dadgar, et al., 2014).

Muscle regeneration involves highly synchronized activations of various cellular processes, and here we ask whether an unsupervised deconvolution of bulk tissue gene expression data is able to discern the molecular subtypes and proportional dynamics. We applied CAM to a time-course gene expression dataset obtained from a mouse muscle regeneration process (GSE469). The time-course gene expression data were acquired over 27 successive time points after the injection of cardiotoxin into the mouse muscle, which induces staged muscle regeneration (**Fig. 1a**). The MDL criterion suggests *K* = 4 as the number of molecularly distinct subtypes indicated by the four vertices of scatter simplex (**Fig. 1d**), and CAM uncovers the temporal repopulation dynamics of the four *de novo* subtypes at each time point (**Fig. 1b**), over 214 subtype-specific markers (**Fig. 1c**). Using gene set enrichment analysis, these subtypes are found to be closely associated with the activation, proliferation, differentiation, and maturation of muscle regeneration.

By a closer look into the subtype-specific gene expression profiles estimated by CAM in a separate study on muscular dystrophies (Dadgar, et al., 2014), we found that TGFβ-centered networks strongly associated with pathological fibrosis and failed regeneration were also induced during normal regeneration, but at distinct time points (temporal parsing into sub-networks). We hypothesized that asynchronously regenerating microenvironments was the underlying driver for fibrosis and failed regeneration. We validated this hypothesis using an experimental model of focal asynchronous bouts of muscle regeneration in wild-type mice. Laser capture microdissection and mRNA profiling of each notexin-injection site, and the in-between area was done, showing reproducibly different genome-wide microenvironment data (Dadgar, et al., 2014). Using CAM based tissue deconvolution results and large human biopsy data sets, we have developed a novel model of the gradual failure of regeneration as a function of age/time in the muscular dystrophies, with experimental evidence in mouse models, where asynchronous remodeling puts microenvironments into a ‘arrested development’, unable to progress normally through the time-dependent regeneration process (Dadgar, et al., 2014). This model will have broad implications for many chronic inflammatory states.

### Human brain lifespan

We first re-validated CAM based deconvolution on the benchmark mouse brain data involving neuron, astrocytes, and oligodendrocytes cell types (GSE19380) (Kuhn, et al., 2011). The individual gene expression profiles of the three primary subtypes were variably and experimentally mixed, where the cells were cultured and the mRNAs were extracted separately. Without using any prior information, CAM accurately detected the number of subtypes, identified subtype-specific markers, and estimated the proportions and subtype-specific gene expression profiles. These results were assessed against the ground truth (Wang, et al., 2016). Importantly, CAM successfully detects not only the majority of the known but also many de novo marker genes associated with brain cell types.

We then applied CAM to the benchmark human brain lifespan data set, Human Brain Transcriptome (HBT) (GSE25219, Illumina Human 49K Oligo array, 923 samples with 17,565 probes) (Kang, et al., 2011). Using the time-courses of the bulk gene expression profiles sampled at various human brain developmental periods and lifespans, CAM first detect *de novo* subtype-specific markers (**Fig. 2b**), i.e., the vertex-residing markers detected by CAM directly from the simplex of bulk gene expression data. These blindly detected markers match well with the *a priori* markers associated with the cell subtypes in the human brain (**Table S1**). Based on the expression patterns of these blindly detected markers, CAM then reconstructs the temporal repopulation dynamics of five molecularly distinct subtypes or cell states that were undetectable by either global profiling or supervised deconvolution (**Fig. 2c**). We also separately applied CAM to region-specific HBT data sets of the bulk tissues sampled from the four different cortex regions. The obtained region-specific temporal repopulation dynamics are similar to those estimated from the whole data set (**Fig. S1**). We further replicated the HBT deconvolution results in an independent benchmark data set, Braincloud (GSE30272, Affymetrix Human Exon 1.0 ST Array, 269 samples with 30,176 probes) (Colantuoni, et al., 2011). The deconvolution result in terms of brain cell repopulation dynamics is consistent with HBT (**Fig. 3**). Again, the blindly detected markers in this replication study match well with the *a priori* markers associated with the cell subtypes in the human brain (**Table S2**).

**Figure 2.**
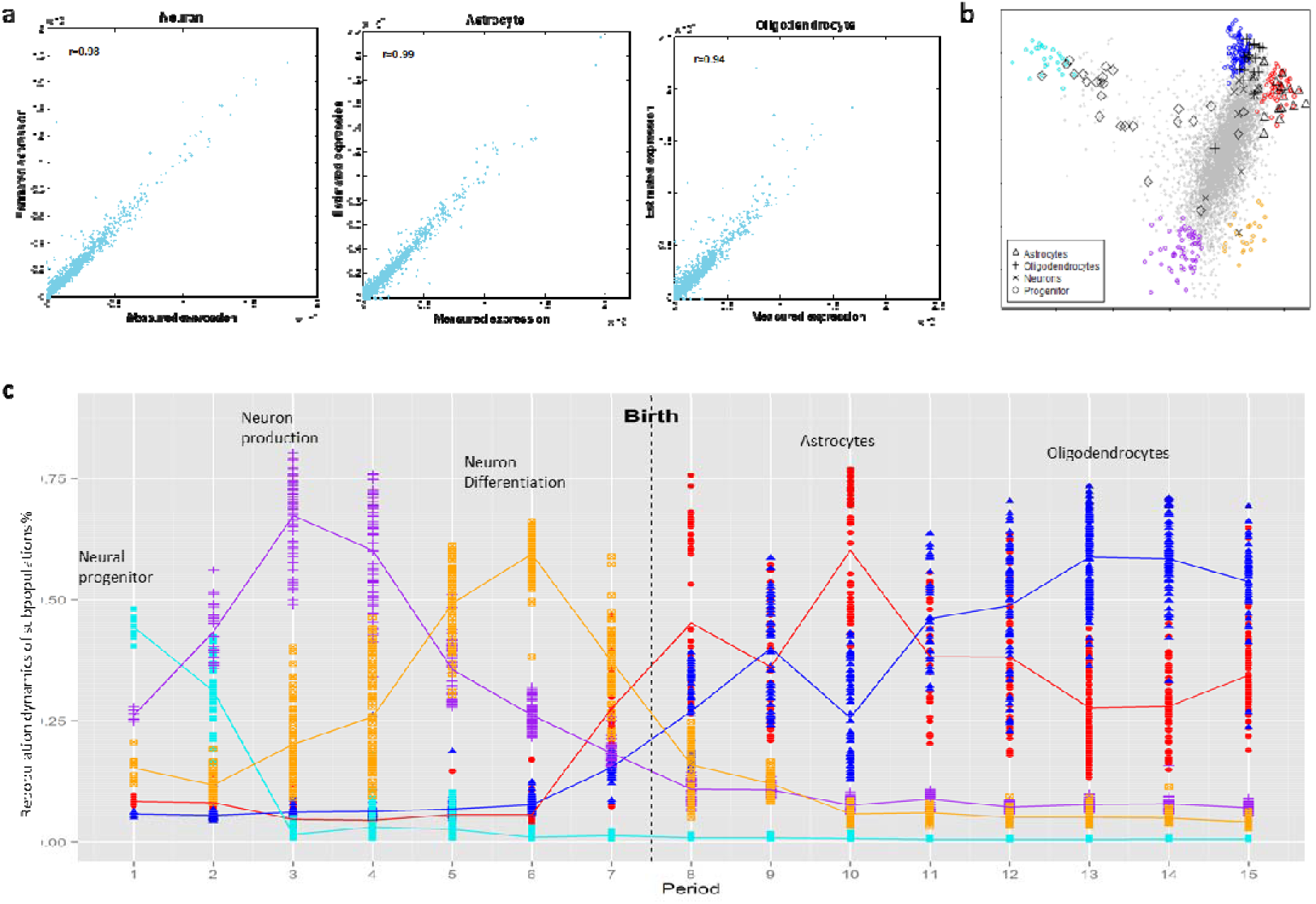
Validation and application of CAM in identifying *de novo* subpopulation-specific marker genes and deconvoluting mixed gene expression profiles, with distinct subpopulations of progenitor, neuron, astrocyte, and oligodendrocyte, at their various development states. (**a**) Scatter plots of the estimated subpopulation-specific gene expression profiles deconvoluted from heterogeneous tissue samples (GSE19380) by CAM, plotted against the measured gene expression profile from pure subpopulations, with an almost perfect correlation coefficient. (**b**) Identification of marker genes by CAM from the simplex of mixed gene expression data (HBT GSE25219) (Gray dots: all genes forming scatter simplex. Light-blue dots: marker genes of neural progenitor. Magenta dots: marker genes of neuron in production. Brown dots: marker genes of neuron in differentiation. Red dots: marker genes of astrocyte. Blue dots: marker genes of oligodendrocyte. Gray symbols: *a priori* marker genes for four known cell subpopulations.) (**c**) Based on the gene expression time-courses of cell subpopulations in human brain sampled at various developmental periods, CAM detected *de novo* subpopulation/state-specific marker genes and dissected mixed gene expression profiles into subpopulation/state-specific gene expression patterns. The result revealed repopulation dynamics of five distinct subpopulations/states that were undetectable by either global profiling or supervised deconvolution. The relative proportions of these subpopulations or states estimated by CAM, plotted as a function of time, match well with the previously-validated cellular repopulation dynamics during brain development.

**Figure 3.**
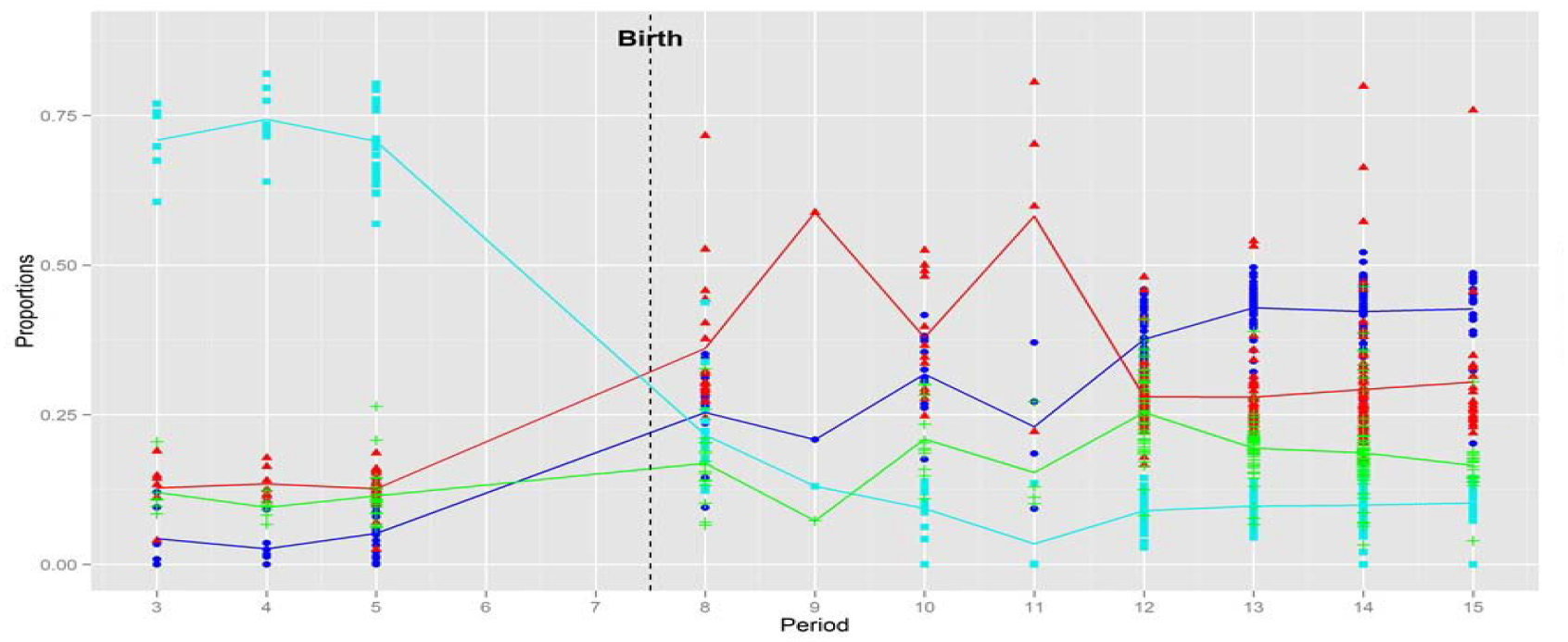
Application of CAM to deconvolve Braincloud data (GSE30272) that replicate the repopulation dynamics of progenitor, neuron, astrocyte, and oligodendrocyte, at their various development states, consistent with what obtained from HBT data.

The relative proportions of these primary brain cell subtypes, blindly estimated by CAM and plotted as a function of time, match well with the previously-validated repopulation dynamics data about human brain development and lifespan (Stiles and Jernigan, 2010). Neuron production in humans begins on embryonic day 42, extends in most brain areas, and is largely complete by midgestation. Among the stem cell lines that emerge during gastrulation are the neural stem cells. The neural stem cells are capable of producing all of the different cells that make up the brain and central nervous system, and for this reason the neural stem cells are usually called the neural progenitor cells (Stiles and Jernigan, 2010). While most neurodevelopmental events involve the proliferation of neural elements, two important processes involve substantial loss of neural elements. These two processes include naturally occurring cell death, which involves the normal loss of 50% or more of the neurons within a brain region; followed by the systematic elimination of up to 50% of between-neuron connections. cascade. Apoptosis has been documented within all of the neuronal and neural progenitor cell compartments in the human brain. Across the cortex, rates of apoptosis within all layers is high, reaching 70% in some regions. Importantly there is strong evidence of high levels of cell death in the neural progenitor population. population. In contrast, proliferation of glial progenitors, while beginning prenatally, continue for an extended protracted period after birth as oligodendrocytes and astrocytes differentiate. In summary, brain development involves overproduction of neurons and glial cells, followed by various regressive events.

### In-house brain bulk tissue data

Brain is made up of hundreds of different cell types. While the major classifications are neuronal cells and glial cells, each of these has many subcategories based on their morphology or functions (Mancarci, et al., 2017). Our in-house brain data were gene expression profiles measured from 129 parietal cortex (PC) tissues and 129 cerebellum (CB) tissues. The major cell types in PC and CB regions are shown in **Table S3**, with *a priori* markers available from the literatures (Xu, et al., 2013). CAM3.0 identified three and four subtypes, respectively. The enrichment of *a priori* markers in each deconvoluted subtype indicates CAM3.0 successfully finds three major brain cell types – astrocyte, mature oligodendrocyte, and neuron – in the human parietal cortex (**Fig. 4a**). Among much more complex compositions in the cerebellum, CAM3.0 also distinguish four major cell types (**Fig. 4b**). CAM3.0-identified markers located in simplex vertices match well with part of *a priori* markers (**Fig. 4, Table S4-S5**). Two simplex scatter plots reflect astrocyte and mature oligodendrocyte, as two major glia cell types, have distinct patterns at molecular expression level from others in both cortex and cerebellum. The CAM3.0-estimated proportions of cell types (**Fig. S2**) indicate neuron cell types account for a larger proportion than the sum of two major glia cell types.

**Figure 4.**
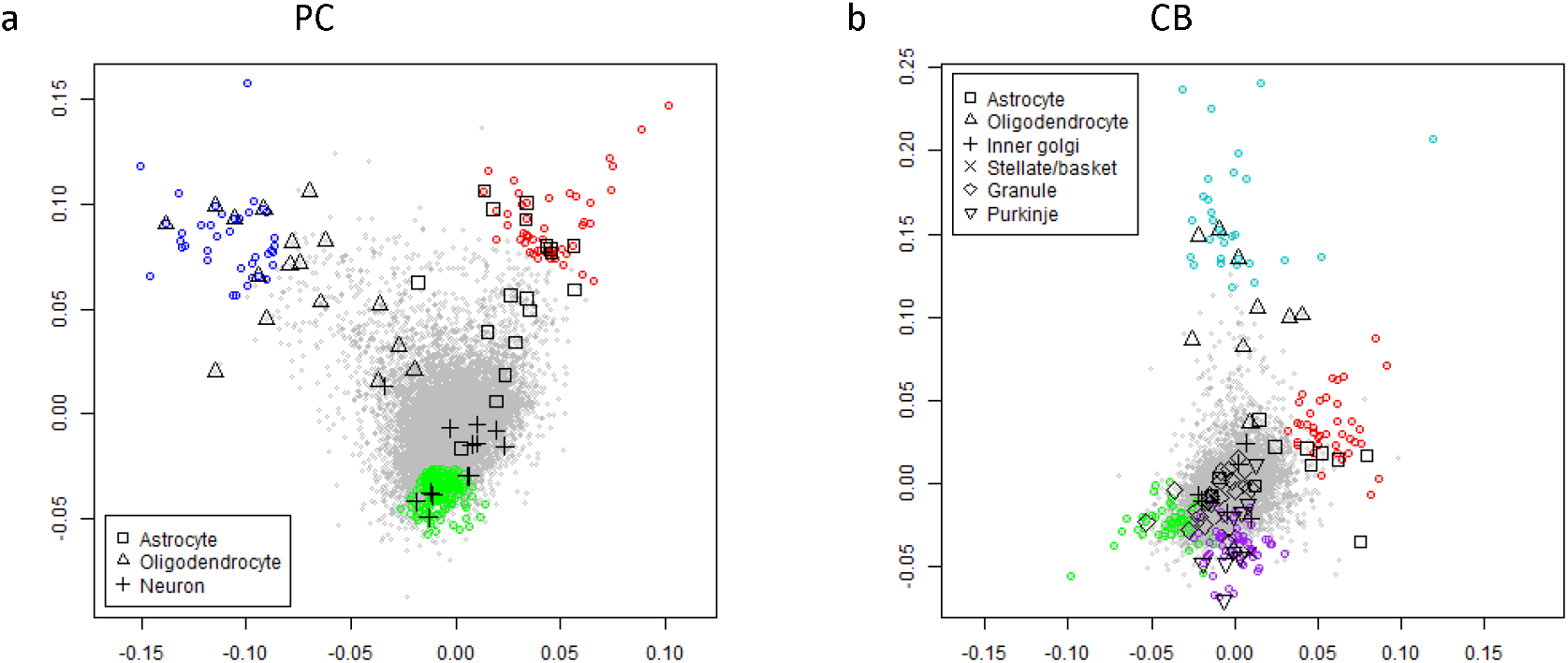
Scatter simplex of complex tissues from parietal cortex and cerebellum (Gray dots - all genes forming scatter simplex; colored dots – markers identified by CAM3.0; black symbols - a priori markers for known cell types)

While the two major types of cells in the brain are known to be glia and neuron, the true ratio of glia to neurons in the brain remains a mystery. One of the recent studies using an efficient cell counting method provides compelling evidence for 1:1 ratio on four whole human brains (Azevedo, et al., 2009). The same study also reveals that the ratio of glia to neurons in the brain varies from one region to another, sometimes dramatically, e.g., 3.76:1 in the cerebral cortex versus 1:4.3 in the cerebellum (Azevedo, et al., 2013; Azevedo, et al., 2009). However, other scientists have argued that more rigorous studies are needed in which just about every known or unknown marker for both neurons and glia is used to capture as many of the different cell types as possible.

### Vascular bulk tissue proteomics data

At the molecular level, atherosclerosis can be defined as an assembly of hundreds of intra- and extra-cellular proteins that jointly alter cellular processes and produce characteristic remodeling of the local vascular environment. Ultimately, these proteomic changes produce the lesions responsible for most ischemic cardiovascular events. The ability to identify marker proteins characteristic of early- and late-stage pathological tissue states would have meaningful clinical impact, but challenged by the heterogeneous nature of whole tissue proteomic profiling involving varying proportions of different tissue phenotypes.

In one of our projects for Global Analysis of Coronary and Abdominal Aortic Proteomes, specimens were collected from left anterior descending coronary artery (LAD) and abdominal aorta (AA) regions in 99 donors free of clinical/diagnosed cardiovascular disease (**Fig. 5**). A trained pathologist scored each specimen for surface involvement of fatty streak (FS), fibrous plaque (FP) and normal (NL). 1583 proteins in LAD and 1273 proteins in AA are quantified by the DIA-MS technique with less than 50% missingness. After data normalization and imputation, CAM3.0 algorithm was performed and the results indicated four and two distinct expression subtypes in the LAD and AA specimen, respectively (**Fig. 6**). We also applied a supervised method called csSAM, which deconvolute **S** from **X** with prior **A**, to estimate tissue subtype-specific expression profiles by pathologist-scored proportions of three tissue subtypes and further obtained associated markers using OVE-FC scheme. Although subtype proportions observed by the pathologist were quite crude, the overlap of CAM3.0-identified markers and csSAM-identified markers (**Table S6** and **Table S7**, numbers in parentheses are counts of markers) provided clues about the biological interpretation of CAM3.0-identified subtypes in LAD/AA. While FS and NL subtype in AA were not separated by CAM3.0 due to insufficient mixture diversity, LAD detected an extra subtype (CAM3.0-NL1), which may reflect complex compositions of normal tissues.

**Figure 5.**
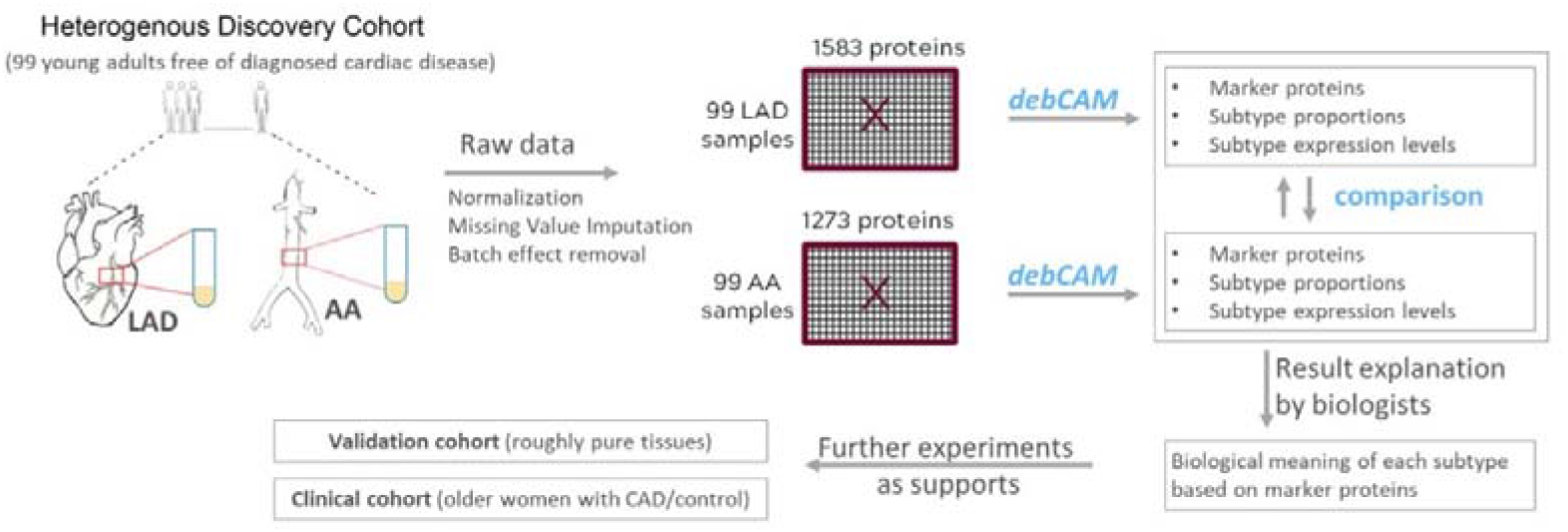
CAM3.0 analysis workflow (implemented in the prototype software debCAM) for proteomics data acquired from two vascular regions (LAD and AA). CAM3.0 results are cross-validated by comparing marker proteins detected in two regions, interpreted by biologists and further supported by biological validation cohort and clinical cohort.

**Figure 6.**
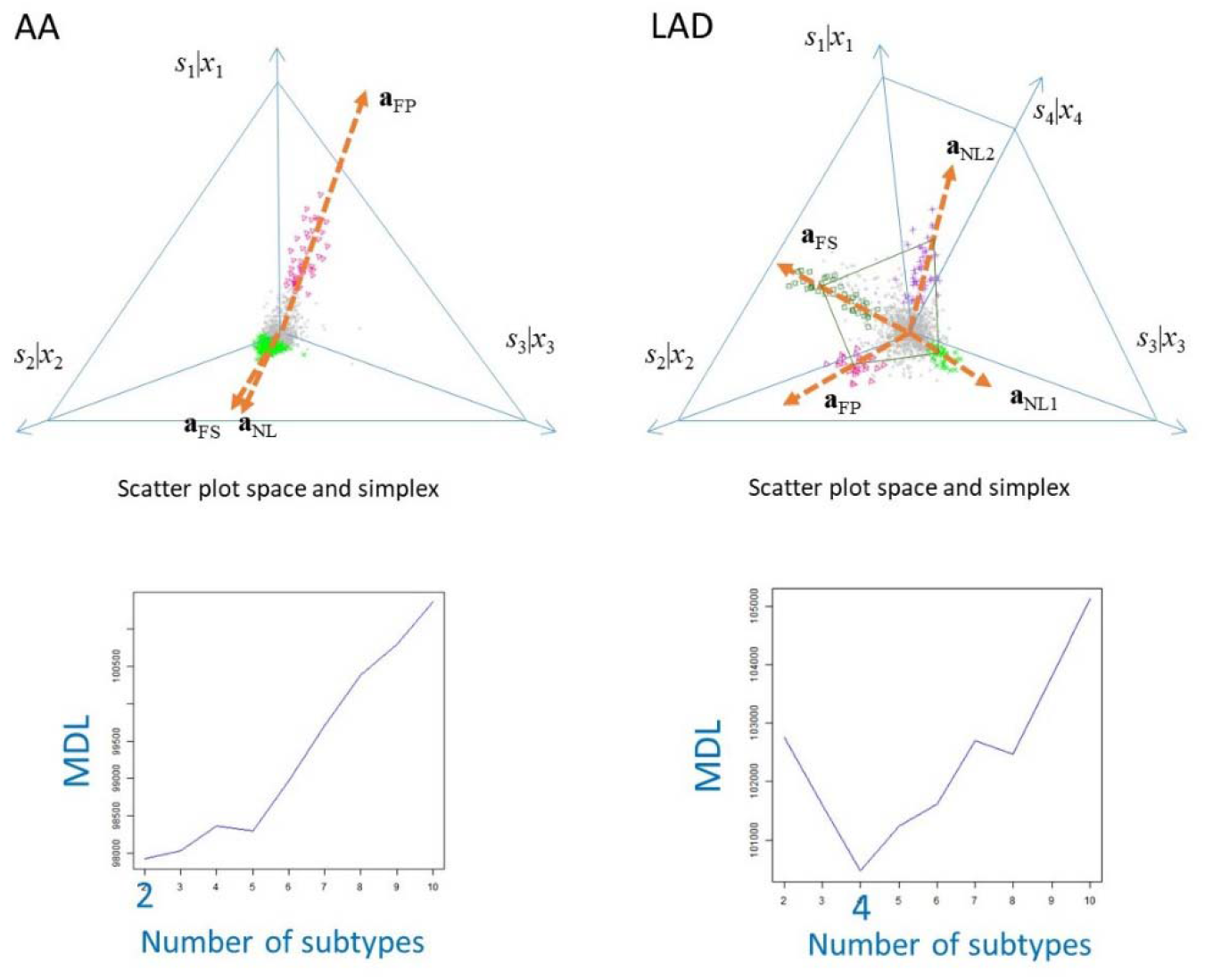
The scatter simplex and MDL curve associated with AA and LAD samples.

As pathologist-scored proportions were just visual inspection of the arterial samples and thus less reliable than CAM3.0 estimations, our collaborator biologists started focusing on CAM3.0-identified subtypes, especially FP subtype whose abundance represents the vascular FP burden and indicates more severe status than NL and FS. Biologists searched the literatures and public database to confirm that CAM3.0-identified FP markers are associated with lesion/FP. Besides, the 34 FP markers blindly detected from LAD and 49 FP markers from AA have an overlap of 21(33.9%), cross-validating the unsupervised discovery from each other.

As the enrichment levels of FP markers will be good candidates for representing the vascular FP burden and marking early atherosclerosis, biologists conduct two more experiments to further support putative FP markers from CAM3.0. The first validation cohort collected a set of specimens isolated from an orthogonal, separate cohort of donors. These pure aortic tissues are isolated from regions with no pathology (pure normal, n=3) and regions completely comprised of fatty streak (pure FS, n=3) or fibrous plaque (pure FP, n=4) luminal surface involvement. 1478 proteins were quantified from the pure specimens by the same DIA-MS technique as LAD/AA tissue profiling used so that expression levels could be comparable from two independent experiments. 58 of 62 putative FP marker proteins are quantified in a validation cohort and most are enriched in pure FP specimens compared to pure FS and pure normal specimens. Statistical comparison (OVESEG-test (Chen, et al., 2019)) of the expression profiles among pure FP, pure FS and pure normal tissue specimen shows 18 of 58 putative markers have a significant enrichment with q-value < 0.02. The expression values of FP-marker estimated from either LAD or AA correlated well with those measured from pure specimens, with correlation coefficients of 0.661~0.919 (**Table S8**). The second validation cohort collected clinical data to examine putative FP markers’ performance in classifying coronary artery disease (CAD) patients and healthy women. High intercorrelation was observed among FP marker proteins. Thus, we applied elastic net variable selection within a logistic regression analysis to select a panel of 10 proteins that achieved high AUC and low misclassification rate (Parker, et al., 2017).

### Comparative evaluation on debCAM and CAM3.0

The first dataset for the comparison is RNA mixtures from the rat tissues (GSE19830) (Shen-Orr, et al., 2010), The mixtures contain rat brain, liver and lung specimens (three tissue types) with different proportions. For each proportion combination, there are three technical replicates, totally 33 samples, and 31,099 RNAs. The source number pre-estimation module (deep-learning) detects three distinct subtypes that matches the ground truth. The number of subtypes is further confirmed by the MDL curves (**Fig. 7**). We evaluated the accuracy of deconvolution outcomes by debCAM and CAM3.0 based on the correlation coefficient and cosine values of estimated subtype proportions and the ground truth (**Table S9** and **Table S10**).

**Figure 7.**
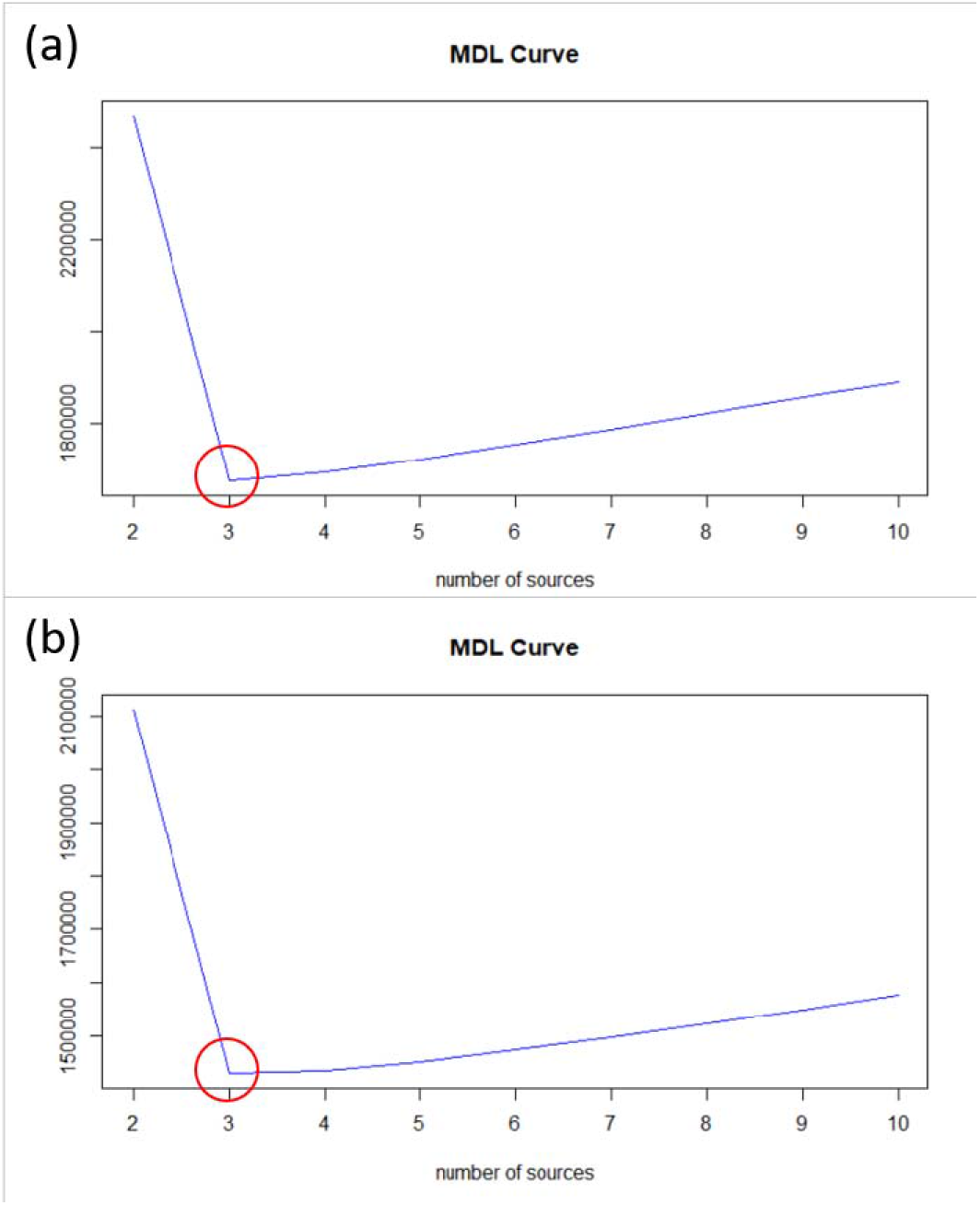
MDL curves on GSE19830. (a) The MDL curve produced by debCAM. (b) The MDL curve produced by CAM3.0. Clearly and consistently, the lowest point of the MDL curves indicates the correct number of subtypes present in the data.

### Application of CAM3.0 to mixtures of immune and cancer cell lines

We evaluate the performance of debCAM and CAM3.0 on a biologically mixed gene expression dataset (GSE64385) (Becht, et al., 2016). In the mixtures, there are five immune cell types and one cancer cell line (six sources). In the dataset, there are 12 samples with different proportions of cell types and 54675 RNAs. The model selection using the combined deep learning and MDL strategies consistently and correctly suggests six subtypes present in the data. Again, we evaluated the accuracy of deconvolution outcomes by debCAM and CAM3.0 based on the correlation coefficient and cosine values of estimated subtype proportions and the ground truth. The results are highly accurate and consistent.

### Re-analysis of braincloud data

We re-analyze the benchmark human brain dataset - Braincloud (GSE30272) using CAM3.0 (Colantuoni, et al., 2011). The samples in this dataset are all human prefrontal cortex of post-mortem brains at different ages, which include fetus, infant, child, teen, adult, and aged. There are 269 samples and 30176 RNAs. The definition at each stage (Kang, et al., 2011) and the corresponding sample number is given in **Table S11**. We first applied deep-learning model to pre-estimate the number of subtypes and obtained a value of six. We then applied MDL criterion to finalize the model selection and observed that the lowest point of the MDL curve is at seven (**Fig. 8**). The subtype-specific markers detected by CAM3.0 are superimposed onto the scatter simplex (**Fig. 9**), and the association with the known primary brain cell types (astrocytes, mature oligodendrocyte, progenitor cell, and neuron) are given in Table 1. From the results, we can see that there seven overlapping markers between the blue source and the progenitor cell, so we can assume that the blue source could be the progenitor cell. Also, since there are 21 overlapping markers, the green source could be oligodendrocyte. Moreover, there are ten overlapping markers, the orange source could be astrocyte. The cyan source is very interesting, since the overlapping markers include astrocytes, oligodendrocyte, and progenitor cell. Thus, one possibility is that the cyan source represents the transition stage of the progenitor cell. That is, the progenitor cell is differentiating. The red and purple sources have only one overlapping markers with neuron, so perhaps they are two different subtypes of neuron. The black source has no overlapping to any prior markers, so it may be a novel unknown subtype, or simply a known cell type but lack of known markers.

**Figure 8.**
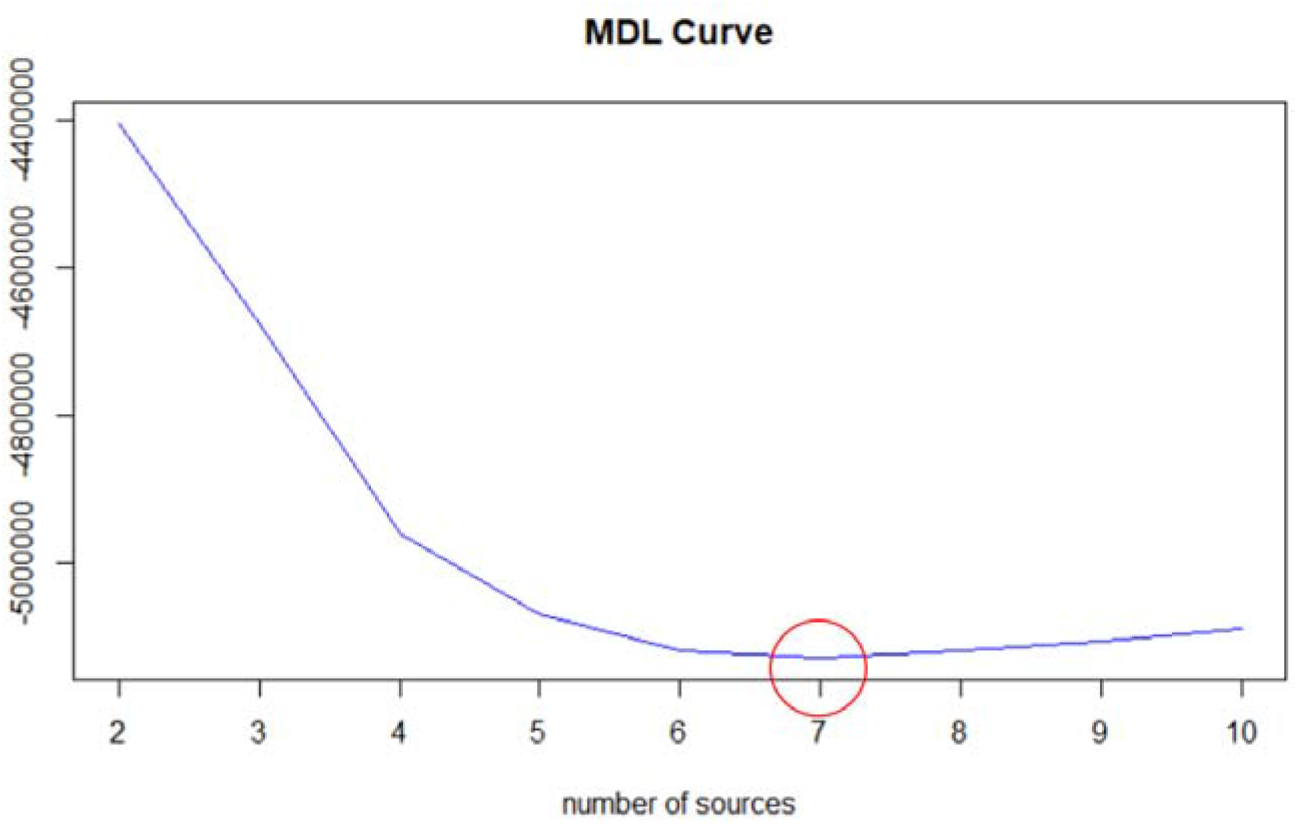
MDL curve for GSE30272. The lowest point of the MDL curve is at seven. Though it is different from the pre-estimation by the deep learning model, they are high consistent.

**Figure 9.**
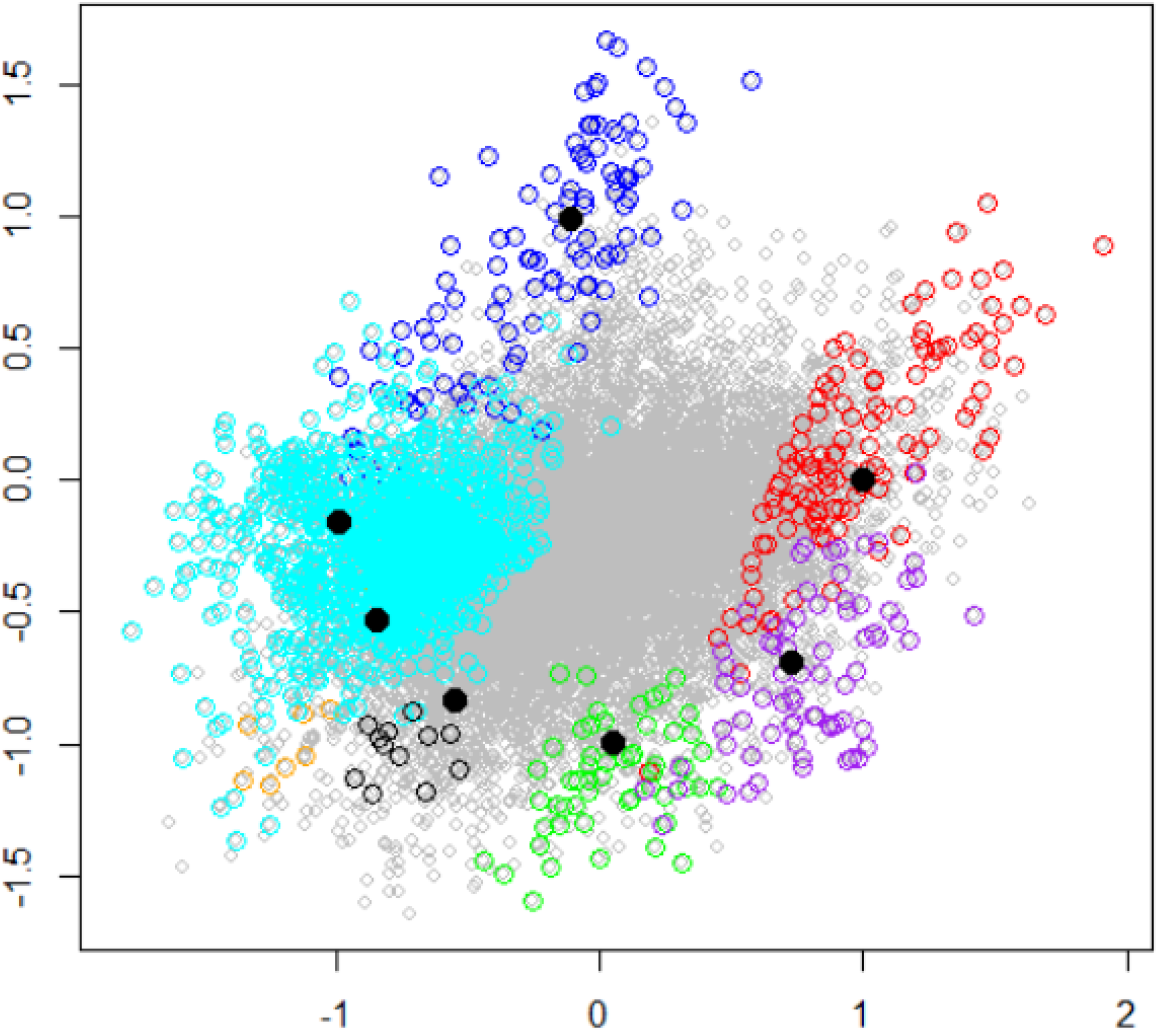
Scatter simplex with superposed color-coded subtype markers. The red, blue, green, orange, purple, cyan, and black circles are the SMGs detected by CAM3.0 (each color corresponding to one subtype), and one can see that they are all located in the vertices of the simplex, which meets the requirements of SMG.

**Table 1.** Association between a priori markers and CAM3.0 markers. From the table, we can know that the blue source could be progenitor cell, the green source could be oligodendrocyte, and the orange one could be astrocytes.

| Overlapping markers | Astrocytes (50) | Oligodendrocyte (31) | Progenitor cell (35) | Neuron (21) |
| --- | --- | --- | --- | --- |
| Red | 0 | 0 | 0 | 1 |
| Blue | 0 | 0 | 7 | 0 |
| Green | 0 | 21 | 0 | 0 |
| Orange | 10 | 0 | 1 | 0 |
| Purple | 0 | 0 | 0 | 1 |
| Cyan | 10 | 1 | 2 | 0 |
| Black | 0 | 0 | 0 | 0 |

As aforementioned, as a time series dataset, we can see the cell type proportion transitions across time after deconvolution. That is, we can plot the proportion matrix with different time period, as shown in **Fig. 10**. The x-axis is the period defined in the **Table S11**, and the y-axis is the proportion of the sources. The points represent the samples in the same period after averaging for each source. From the figure, we can see that the blue source (progenitor cell) decreases significantly after birth, which is reasonable since most of the progenitor cells differentiate during the fetus stage (Ferenczy, et al., 2013). Thus, the increasing of green (oligodendrocyte), orange (astrocyte), and red (neuron) sources is also expected. The cyan source is always in low proportion, the reason may be that the differentiating progenitor cell would not be too many at the same time. Nevertheless, the proportion of the black source is always no lower than 15%. Thus, this unknown cell type in human brain deserves further investigation. In conclusion, the CAM3.0 successfully identified several meaningful sources which are corresponding to known subtypes, and identified some novel subtypes. Also, the deconvolution result of CAM3.0 may help us understand the development of human brain at different period.

**Figure 10.**
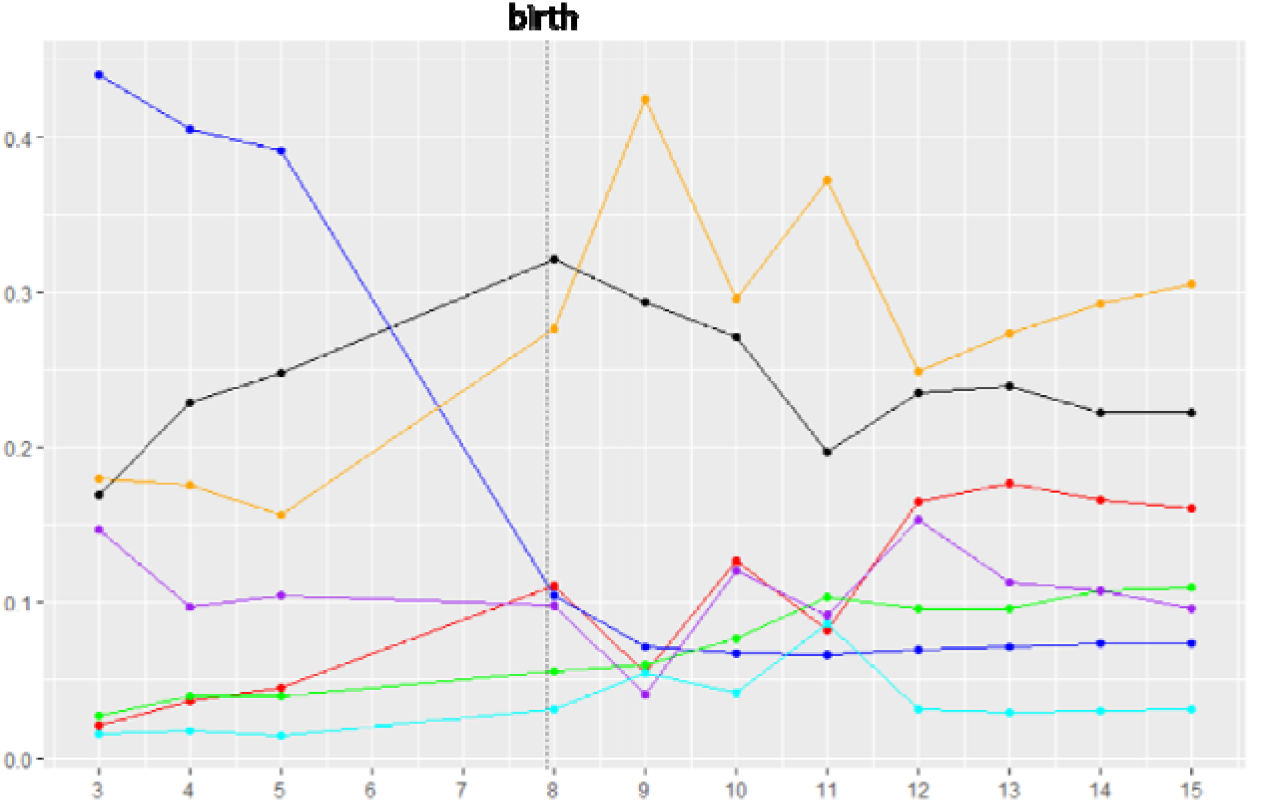
Proportion of each source (A matrix) at different period. The points represent the samples in the same period after averaging for each source. The blue source (progenitor cell) decreases significantly after birth. The green (oligodendrocyte), orange (astrocyte), and red (neuron) sources is increasing after birth. The cyan source is always in low proportion, the reason may be that the differentiating progenitor cell would not be too many at the same time. Nevertheless, the proportion of the black source is always no lower than 15%. Thus, this unknown cell type in human brain deserves further investigation.

### Preliminary application of CAM3.0 prototype to metabolomics data

Recent advances in high-throughput mass-spectrometry (MS) make it possible to perform highly accurate, precise, and sensitive metabolomic profiling on thousands of biologic samples. Unlike conventional targeted metabolomics, un-targeted metabolomics uses an MS assays to access a broader range of metabolites (both known and unknown) than possible from any single metabolomic assay or target list. These un-targeted metabolomic profiles provide a powerful and unbiased means to document the metabolic consequences of behaviors, environmental exposures and risk factors, including genetic risk factors, which influence health and disease.

Using previously obtained un-targeted metabolomic data from subsets of people enrolled in the Multi-Ethnic Study of Atherosclerosis (MESA, N=4,000), we apply CAM3.0 prototype to identify metabolomic archetypes and their relative contributions to individual samples represented by the MS features. The major data acquisition and preprocessing protocols are outlined in **Figure S3**. In the preliminary experimental results, the MDL curve indicates about 9 metabolomic archetypes associated with the CH1 subset of the MESA cohort (**Fig. 15a**), and the heatmaps of marker-enriched archetype expressions show clear archetype-specific patterns (**Fig. 15b**) where the refinement and rearrangement of marker selection by COT improves the quality of the initial markers detected by CAM3.0 prototype directly from the metabolite mixtures.

**Figure 11.**
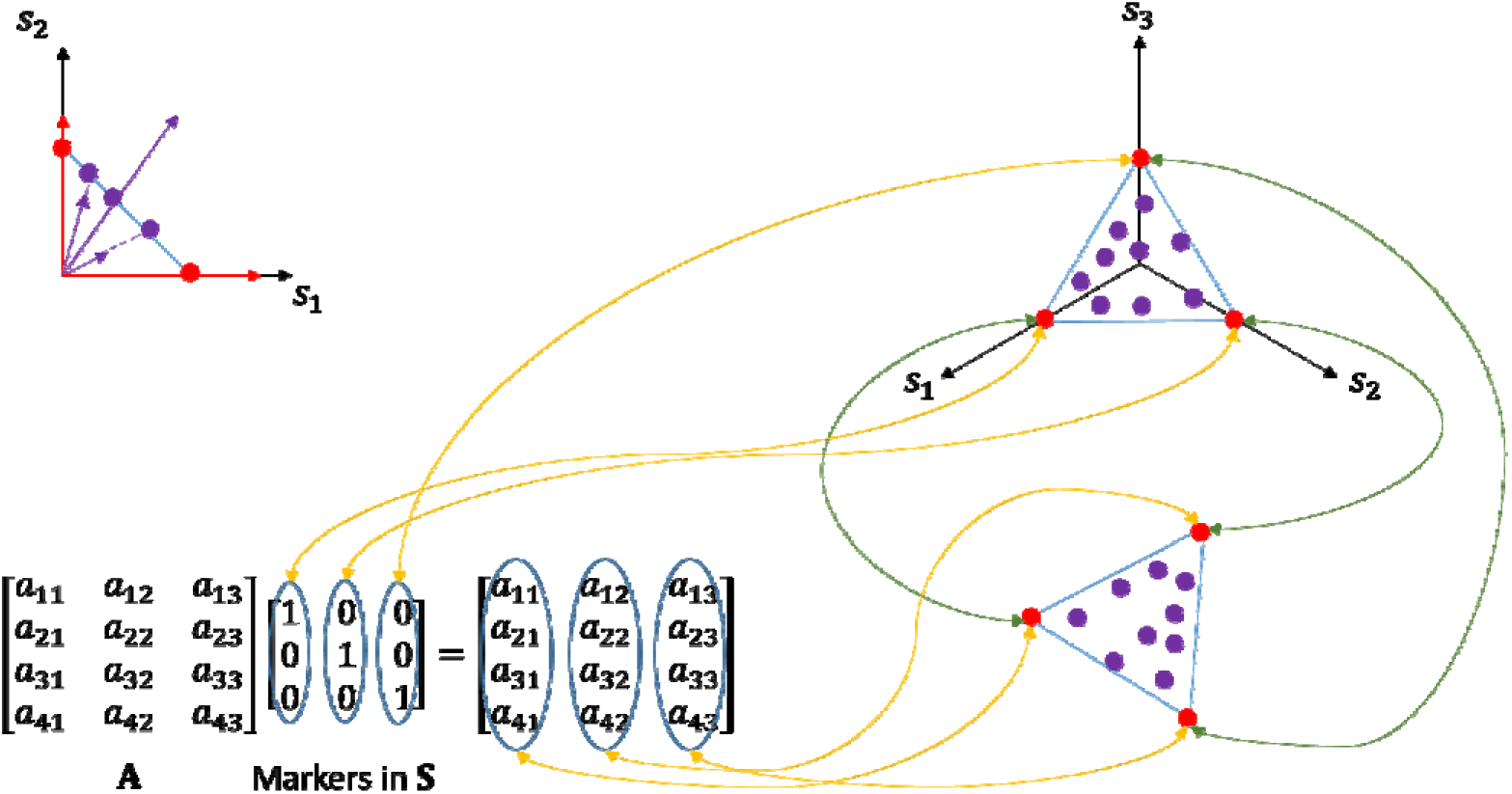
Illustration of the relation between S and X (K = 3 for example). The upper left corner is an example of sum-to-1 normalization in 2D space, which project the genes onto 2 1-simplex (a line). The reds are the marker genes, and the purples are the normal genes. The right part is the simplex in 3D space (a triangle). After multiplying with A, the simplex is rotated and projected into a higher dimension space (four in this figure), and the vertices (markers) become the column vector of A, as shown in the figure.

**Figure 12.**
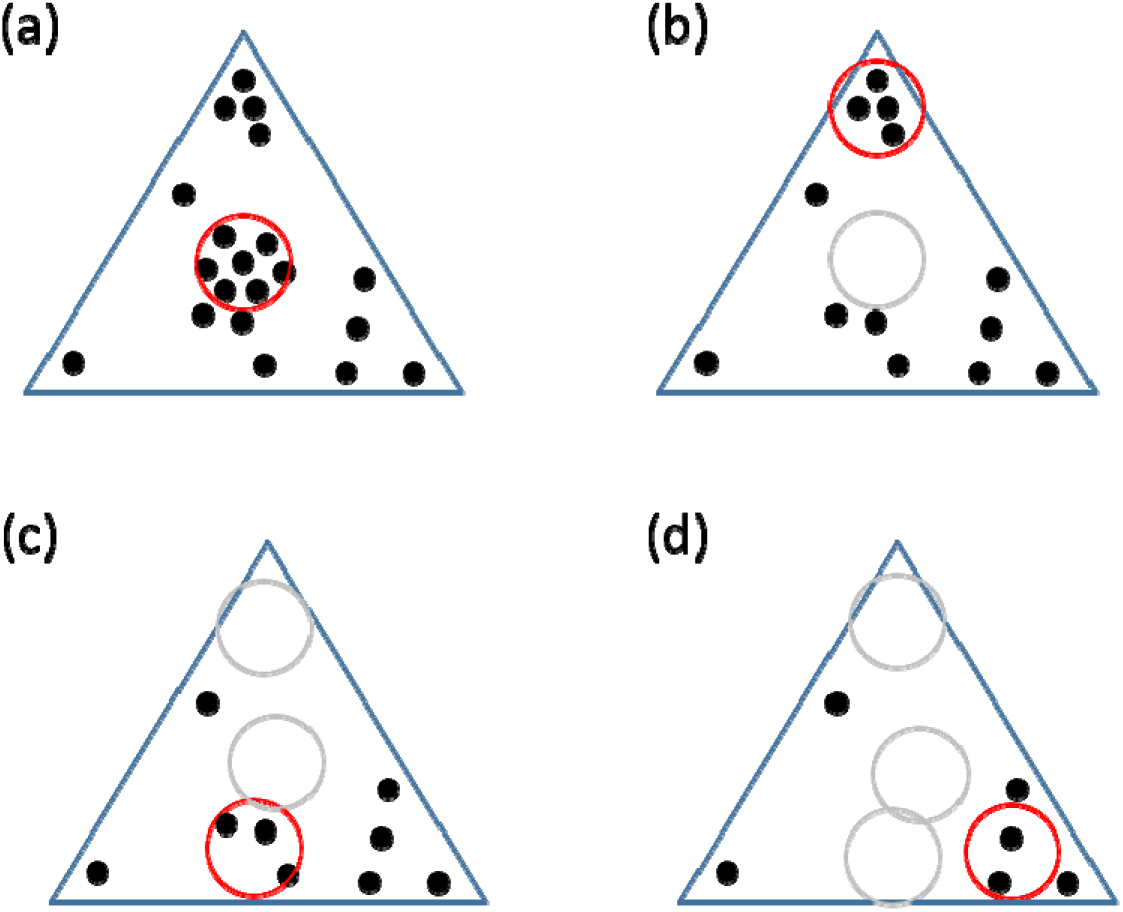
Illustration of radius-fixed clustering. (a) After deciding the radius (the radius of the red circle), we can set each gene as center, and then find which circle contains the greatest number of gene (red circle, seven points). (b) Since the points in the red circle in (a) are removed, the next one contains four. (c) The next one contains three points. (d) The final one contains two, if we set all the clusters should contain at least two points.

**Figure 13.**
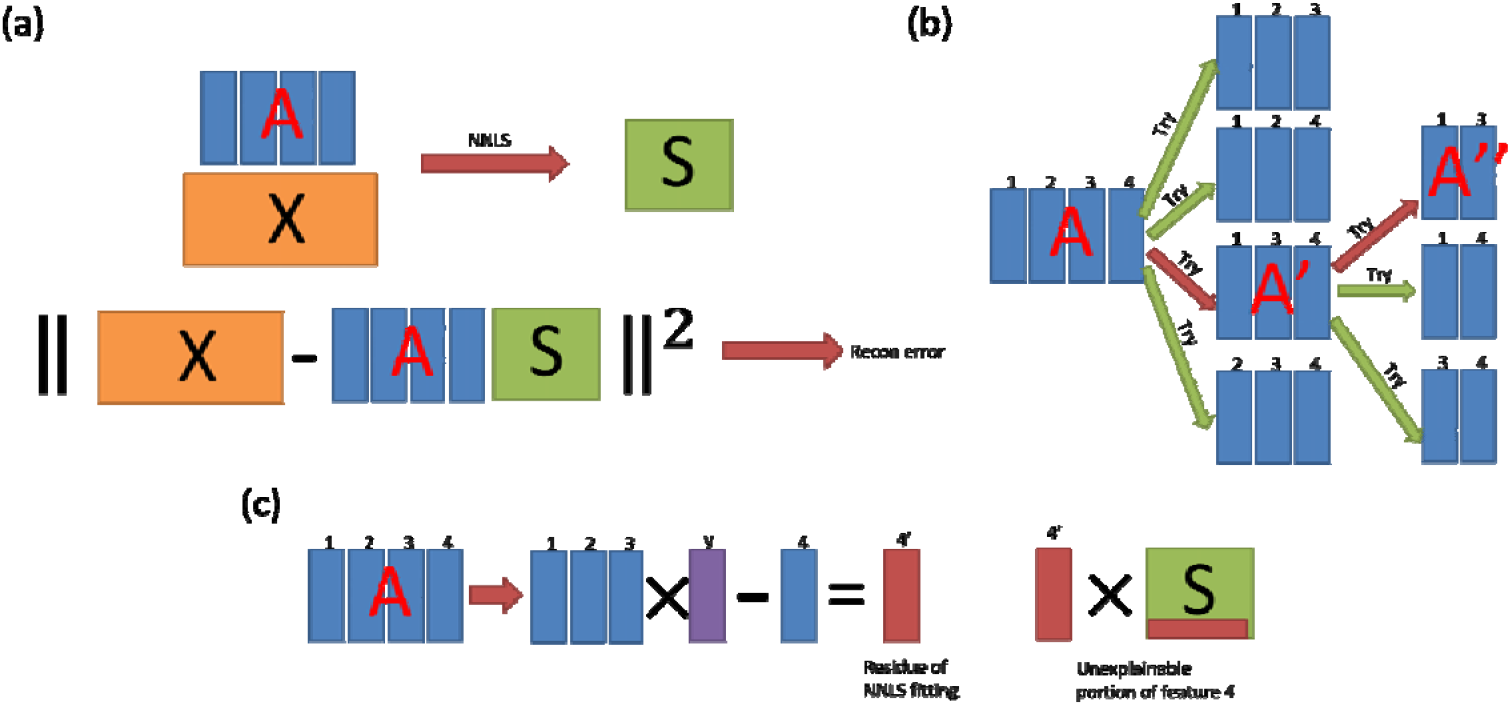
Illustration of SBFS with reconstruction error and unexplainable portion with four features in the feature set. (a) The reconstruction error the sum of the square of the residue by NNLS fitting. (b) In the Step 2 of SBFS, NNLS for fitting ***X*** is performed for excluding each feature in the feature set (four times in this figure) to compute the reconstruction error. (c) Though we still need to perform NNLS four times, but we just need to fit a matrix with only one column, not the whole ***X***, so the computation time decreases significantly.

**Figure 14.**
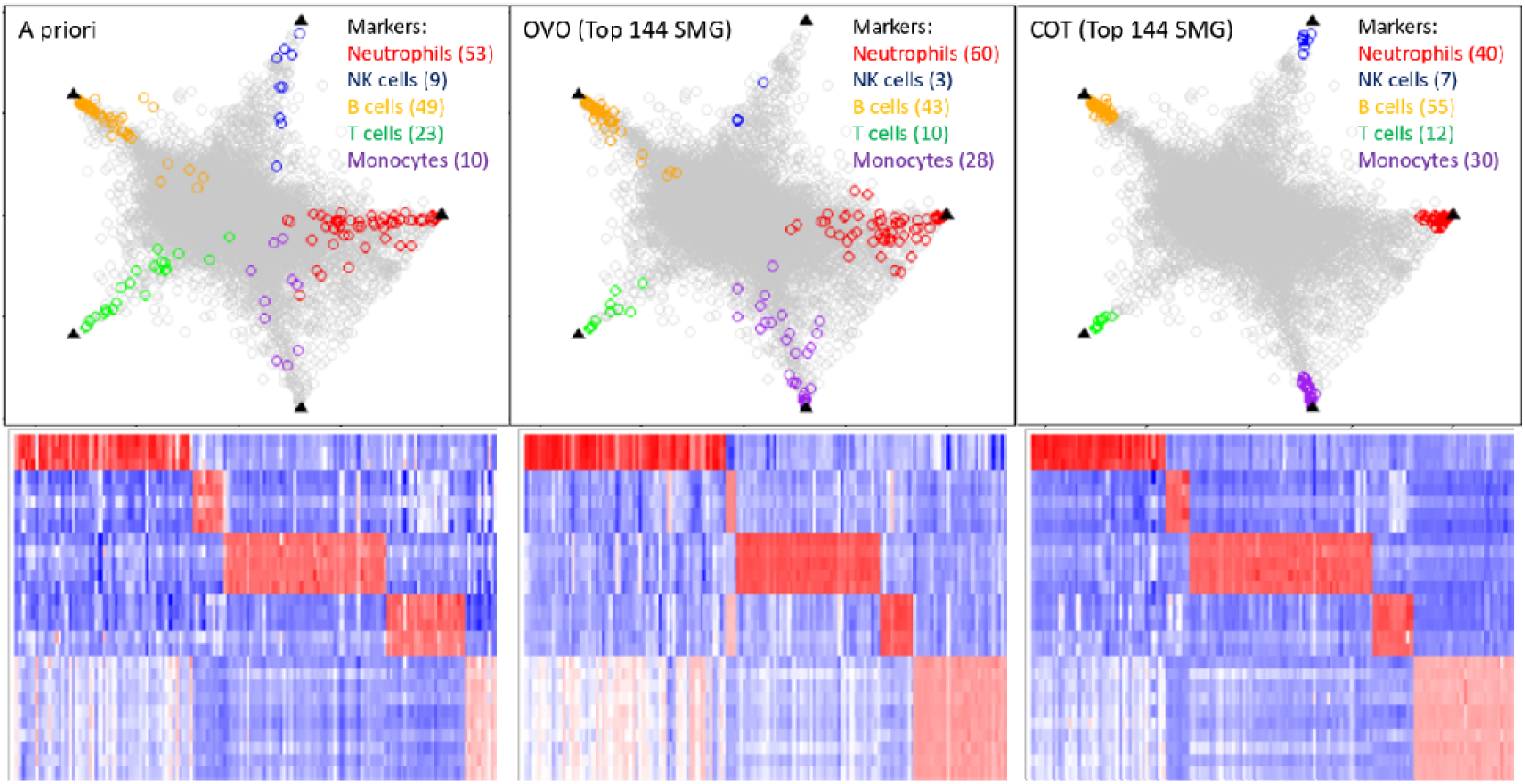
Simplex plots and heatmaps of GSE28490. The black triangles are the exact positions of the vertices of the simplex. In prior knowledge, there are 144 markers in these five subtypes (neutrophils, NK cells, B cells, T cells, and monocytes). The markers detected by a priori and OVO are close to the vertices, but not close enough. However, the markers detected COT are all confined around the vertices. Moreover, in the heatmap of a priori and OVO, it is clear that there some reds (high expression) across different subtypes for some marker genes. In the heatmap of COT, it is clear that except the corresponding subtype, the expressions of the marker genes are all almost blues or whites (low expression).

**Figure 15.**
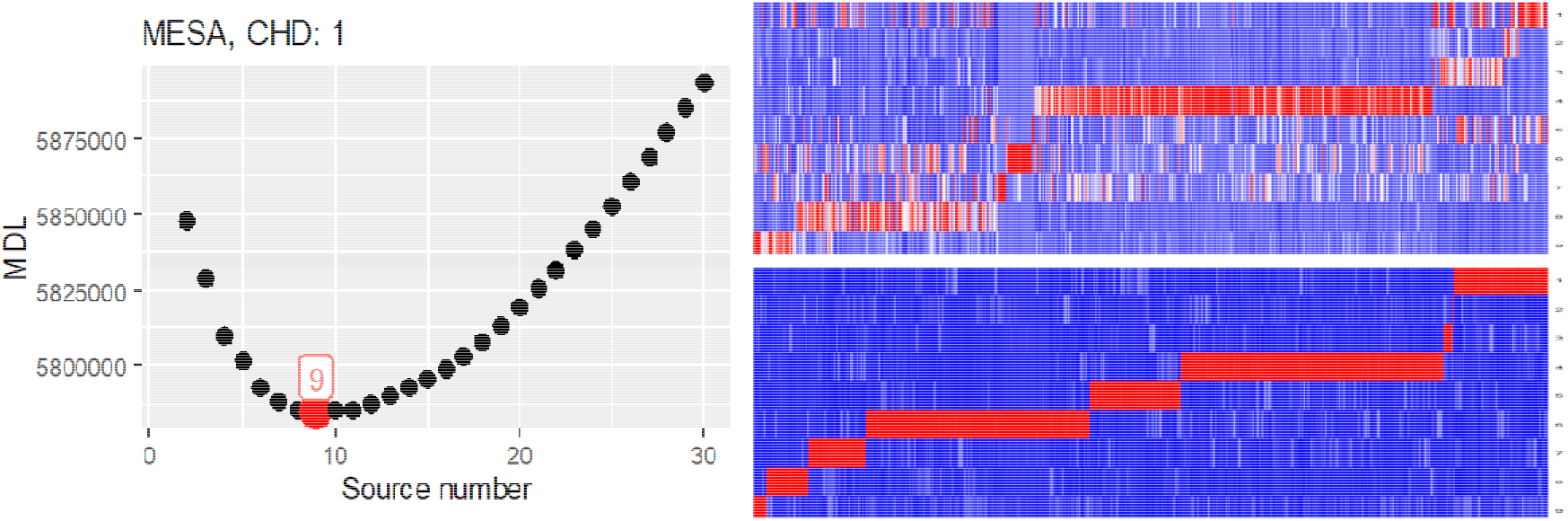
Preliminary results of CAM3.0 prototype application to analyze metabolomics ata.

## 3 Methods

### Mathematical modeling of latent variables

Here we formulate the deconvolution task as a blind source separation (BSS) problem. Supposing the number of samples (observations) is *M*, number of genes (features) is *N*, and number of cell types (sources or subtypes) is *K*, with several assumptions for the application on the tissue heterogeneity deconvolution, the problem can be stated

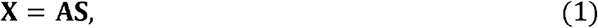

as where **X** is a *M* × *N* gene expression matrix, where each row corresponding to a sample (observation), and each column corresponding to a gene (feature). **A** is a *M* × *K* mixing matrix, where each row corresponding to a sample, and each column corresponding to a molecular subtype (source). That is, each row in **A** represents the portion of each cell type in a sample. **S** is a *K* × *N* source matrix, where each row corresponding to a cell type, and each column corresponding to a gene. Thus, each row in **S** represents the expression in a cell type. Our target is to find out **A** and **S** separately. The only information we have is the observation matrix **X**, so this is a blind source separation problem.

We can assume that for each dimension (each subtype), there is at least one gene as the Cartesian vector (marker gene), though the length is not one. Thus, if we normalize all the genes by their expression sum (the column sums are all 1 after normalization), all the genes are the convex combinations of the marker genes from each subtype. That is, the normalization is corresponding to project the genes onto a hyper plane, where the genes form a (*K*-1)-simplex, and the marker genes are on the vertices of the simplex. Thus, if we multiply **A** to **S**, which is similar to rotate and project the (*K*-1)-simplex into a higher dimensional space, and the vertices of the simplex are the column vectors of **A**. That is, after projection, the column vectors (genes) in **X** can be viewed as convex combination of the column vectors in **A**, so the vertices of the simplex are exactly the vectors in **A**. Thus, after projecting the column vectors (genes) of **X**, we just need an algorithm, such as Quickhull (Barber, et al., 1996), to find the convex hull, and points on the convex hull are the candidates of the vertices. The relation of simplex of **S** and **X** is illustrated in **Fig. 11**.

CAM applies Minimum Description Length (MDL) (Wax and Kailath, 1985) to find the optimal *K*. MDL is a widely-accepted method, which consider the idea of information theory. In general, the idea of MDL is to find the balance between low reconstruction error and low model complexity. The formula of MDL adopted by CAM is:

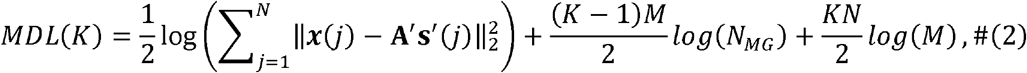

where the first term corresponds to the model fitting error (reconstruction error), and the second and the third terms corresponding to the model complexity. That is, MDL is trying to find a model (*K*) where the reconstruction error is low (first term), and the number of sources (*K*) is also low (second and third terms). We can also view the second and the third terms as penalty terms, which can avoid overfitting problem. With MDL, unlike nICA or NMF, CAM is a fully unsupervised deconvolution method, since the value of *K* could be estimated by CAM automatically, which is not true for the other methods.

The major unresolved problems associated with CAM1.0/2.0 include: (1) unsatisfactory quality of feature clusters specifically related to marker gene clusters, (2) high computational complexity in finding convex full, and (3) high computational complexity in identifying the optimal simplex.

### Radius-fixed feature clustering

While CAM provides two clustering methods in the package, K-means clustering is usually selected. The reason is that when data contains thousands of genes (features) with dozens of samples, the computation time of APC is not feasible. However, as we know, there are some disadvantages of K-means clustering. For example, K-means clustering needs initialization, and which does not guarantee global optimum (MacQueen, 1967). That is, if K-means clustering may give different results with different initializations. Since the clusters from K-means clustering are for simplex vertex identification in CAM, CAM may produce unstable deconvolution results. Also, the reason CAM needs a clustering method for pre-processing is to suppress the noise before finding the convex hull. However, since we cannot control the size of each cluster from K-means clustering, the noise may be suppressed at varying degrees in different clusters, which may distort the shape of the convex hull. Though we can change the number of clusters to affect the size of clusters in K-means, it is not guarantee that the size of all clusters will change with the number of clusters, and the size of all cluster may be still not the same usually. Also, it is well-known that K-means clustering is sensitive to the outliers, so if there are many outliers, K-means clustering is not an ideal choice.

To address the problems of K-means clustering, we want a method which can control the size of each cluster no more than a certain threshold, and the threshold can be control by a parameter easily. Also, it should be a deterministic method so that there is no randomness in the result. Moreover, it would be great that if this method is not sensitive to the outliers. Here we propose another way for clustering, called Radius-fixed clustering. The idea of radius-fixed clustering is that first we need to decide the value of the threshold, then remove the clusters satisfying some requirements one by one from the dataset until there is no clusters which meet the requirements. The threshold here we selected is cosine similarity, since Euclidian distance is distorted after projection, but cosine similarity will not change. The steps of radius-fixed clustering are as following and also illustrated in **Fig. 12**:

1. Set a radius (though we choose cosine similarity for CAM, in other applications, any other kinds of distance or similarity are acceptable).
2. For each gene (point), compute how many genes are within the radius. That is, set each gene as the center of a hyper ball, the radius of which is the one set in step 1, and then compute how many genes are within the hyper ball.
3. Find out the hyper ball with the greatest number of genes inside, and then remove all the genes in the hyper ball from the data.
4. Repeat Step 2 and 3 until there is no gene left or the number of genes in the hyper balls are below a certain value.
5. All the hyper balls removed from the data are the clustering results by radius-fixed clustering.

Since we have set the “radius” of each cluster at the beginning, it is guaranteed that no cluster would be larger than the radius. Thus, we can control the size of each cluster efficiently, or at least we can make sure that when computing the mean of each cluster, there is no gene which is far from the center of the cluster. Also, if we set a threshold for the number of genes in each cluster, there may be some genes which do not belong to any cluster. Since the target of the clustering step is to suppress the noise, it is acceptable that if there are some genes which are not clustered. That is, the target of radius-fixed clustering is to “down-sampling” the simplex for de-noising, not for “clustering” all the genes to clusters. Also, if these “not clustered” genes are outliers, actually it would even make our de-noising more accurate. Moreover, in biology, genes always work together (pathways) (Kelley and Ideker, 2005). That is, a bunch of genes will have similar patterns since they have related functions. Thus, when using radius-fixed clustering, setting a threshold for member number of a cluster can not only remove the outliers, but also follow the biology principle.

### Convex hull identification

The core idea of CAM is to identify the marker genes, which are located on the vertices of the simplex. However, it is hard to find the optimal simplex directly, so CAM finds the convex hull of the data first. That is, the optimal simplex should be a subset of such complete convex hull. Currently, the most popular method is Quickhull (Barber, et al., 1996), a divide and conquer approach, the idea of which is similar to quicksort. Basically, Quickhull is trying to find a convex set and remove the internal points iteratively, until no points can be removed. With respect to the number *N* of features for 2 or three dimensions, the computational complexity of Quickhull is about O(*N* log *N*). For a *d*-dimensional case (i.e., *d* samples or mixtures), the computational complexity becomes 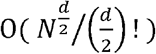.

Recall that the objective is to find the extreme points in the data. By the definition of convex set, any point *p_i_*, in the data can be the convex combination of all points:

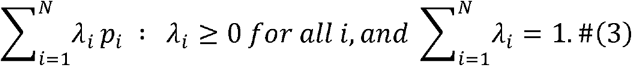

However, the definition of the extreme point is that the point can be constructed by just itself (the coefficients of the other points are all zero). That is, we are finding some points the linear combination of which are only themselves with some constraints on the coefficients. This problem can be formulated as a linear programming problem (Pardalos, et al., 1995),

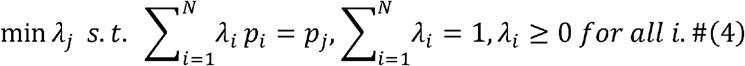

We can test all *P_j_* in the equation 4.6, and if *λ_j_* is not zero, then the corresponding *p_j_*, is an extreme point. There are several advantages if we use linear programming to find the convex hull. First, though the time complexity of linear programming is just weak polynomial, is still faster than the one of Quickhull regarding dimension. Second, the number of clusters in CAM is usually around 100, and all clusters can be tested simultaneously. That is, for linear programming, we can even accelerate the computation with parallel computing. Third, if we “relax” the convex hull, for example, the points near the boundary could be also identified as the extreme points, we can manipulate the constraints of the linear programming.

### Identification of the optimal simplex

Given the number *K* of sources, identification of the optimal simplex from the convex hull is a combinatorial search problem. That is, there is 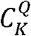 combinations to form a (*K*-1)-simplex. To reduce the computational complexity of this combinatorial search, we propose to directly minimize the reconstruction error via the strategy of combined Sequential Forward Floating Search (SFFS) and Sequential Backward Floating Search (SBFS) (Pudil, et al., 1994). More importantly, in such a greedy search, SFFS or SBFS searches different values of *K* and the optimal simplex, simultaneously, reducing the computational time significantly. The detailed steps of SFFS are as follows (SBFS can be similarly designed):

1. Test all *Q* candidates, and select the one with lowest reconstruction error with **X** by NNLS. Then move the selected one from the candidates to the feature set.
2. Select one from the remaining candidates, and use this one and the feature set to compute the reconstruction error. After testing everyone in the remaining candidates, move the one with the lowest reconstruction error to the feature set. Record the reconstruction error for the current number of features in the feature set.
3. Remove one from the feature set temporarily, compute the reconstruction error for the current feature set, and then move the one back. After testing everyone in the feature set, if the lowest reconstruction error is lower than the recorded for the current number minus one of features in the feature set, remove the corresponding feature and replace the recorded reconstruction error.
4. Repeat Step 3 until no lower reconstruction error can be found.
5. Repeat Step 2 – 4 until there is no candidate left.

Because both SFFS and SBFS will test all possible values of *K* within their loops, we can compare the reconstruction error from SFFS and SBFS for the same source number, and pick the one with lower reconstruction error as the final result for certain source number. The illustration of SBFS is shown in **Fig. 13**.

To further reduce the computational complexity, we also approximate the reconstruction error by the one with the lowest “unexplainable portion” to the others when deciding which candidates should be removed from the feature set. The unexplainable portion of ***X*** is the residue multiplying the corresponding row in the S matrix for each of the feature set fitted via NNLS. Estimating unexplainable portions is efficient because only the selected feature of the feature set needs to be fitted, not the entire ***X***.

### Detection of subtype-specific markers

Ideally, a molecularly distinct subtype would be composed of molecular features that are expressed uniquely in the cell or tissue subtype of interest but in no others – so called subtype-specific marker genes (SMG). SMG plays a critical role in the characterization or classification of tissue or cell subtypes. We and others have recognized that the test statistics used by most existing methods do not exactly satisfy the SMG definition and often identify inaccurate or unreliable SMG. With the availability of subtype-enriched molecular expression profiles, data-driven software tools to detect SMG accurately are essential to support the subsequent steps in systems biology research. We develop an accurate and efficient method to detect SMG among many subtypes using subtype-enriched expression data. Formulated as a Cosine based One-sample Test (COT) in scatter space, the COT framework is fundamentally different from peer methods in that the newly-proposed test statistic precisely matches the mathematical definition of an ideal SMG. Importantly, COT uses the cosine similarity between a molecule’s cross-subtype expression pattern and the exact mathematical definition of an ideal SMG as the test statistic, and formulates the detection problem as a one-sample test. Under the assumption that most genes are associated with the null hypothesis, COT approximates the empirical null distribution with a finite normal mixture distribution for calculating p-values (Efron, 2004).

The most frequently used methods rely on an ANOVA model where the null hypothesis states that samples in all subtypes are drawn from the same population. Another population method is the One-Versus-Rest Fold Change test or One-Versus-Rest t-test (OVR-FC/t-test) that is based on the ratio of the average expression in a particular subtype to the averaged expression in all other (rest) samples (Chikina, et al., 2015). Alternative strategies include One-Versus-One (OVO) t-test and Multiple Comparisons with the Best (MCB) (Newman, et al., 2015). Importantly, we and others have recognized that the test statistics used by most existing methods do not satisfy exactly the SMG definition and often *ad hoc* OVE set intersections have been used to finalize SMG (Chen, et al., 2021; Patrick, et al., 2020; Wang, et al., 2016).

Mathematically, an SMG *i_SMG,k_* associated with subtype *k* is defined as a gene expressed significantly high in subtype *k* while universally low in any other subtypes, approximately

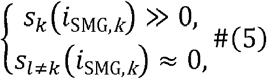

where *s_k_*(*i_SMG,k_*) and *s*_*l*≠*k*_(*i_SMG,k_*) are the expressions of gene *i* in subtypes *k* and *l*, respectively, and are assumed to be nonnegative. Accordingly, the cross-subtype expression pattern of an SMG can be concisely represented by the Cartesian unit vectors 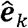, serving as the ground reference for scoring *de novo* SMGs. Fundamental to the success of proposed method is the scale-invariant score cos(***s***(*i*),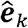) that measures directly the similarity between cross-subtype expression pattern ***s***(*i*) of gene *i* and SMG expression pattern 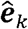 of subtype *k* in scatter space. Specifically, for gene *i* and subtype *k*, the SMG score is given by

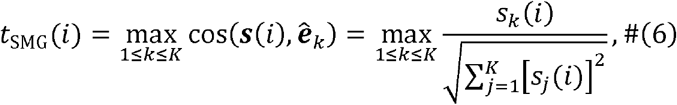

where ***s***(*i*) = [*s*_1_, *s*_2_, …,*s_K_*(*i*)] is the averaged cross-subtype expression pattern of gene *i* over samples, and *K* is the number of participating subtypes. Because ***s***(*i*) is confined within the first quadrant where the central vector is the ‘all-ones’ vector 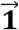, we have 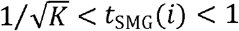. Note that the ‘max’ operation in (6) is specifically applied to handle multiple subtypes.

Following normalization and antilogarithm transformation, the COT software tool performs the following major analytics steps (Figure 1):

1. Data Preprocessing. Molecule features whose norms of cross-subtype expression levels are lower or higher than pre-fixed thresholds are removed as noise or outliers.
2. Test Statistic Calculation. For each of the remaining genes, the cosine similarity cos(***s***(*i*),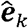) between averaged cross-subtype expression patterns and an ideal SMG reference is calculated.
3. Null Distribution Estimation. The empirical null distribution is summarized over all genes and approximated by a mixture of normal distributions.
4. SMG Detection. Based on the observed test statistic and null distribution, an SMG is identified and subjected to a proper one-sided significance threshold.

We evaluate the performance of proposed method on a benchmark dataset. In the GSE28490 dataset (Allantaz, et al., 2012), we picked out five subtypes with prior knowledge 144 markers for the assessment (Becht, et al., 2016). The geometric proximity of the 144 SMG detected by a priori, OVO, and COT, to the vertices of scatter simplex and the heatmaps are given in **Fig. 14**. It is clear that the markers detected by all three methods are close to the vertices of the simplex. However, for a priori and OVO, it is obvious that the detected markers are not exactly located at the vertices. On the other hand, the markers detected by COT are all confined around the vertices. Moreover, the heatmap of COT shows the markers detected by COT expressed significantly only in the corresponding subtype, but which is not true for a priori and OVO. As shown in the heatmaps, there are some reds (highly expressed) in more than one subtype in a priori and OVO t-test. Given the definition of marker genes, **Fig. 14** presents an example that sometimes prior knowledge may not be accurate enough.

## 4 Discussion

In this report, several methodological improvements are proposed, evaluated and applied to perform full unsupervised deep deconvolution. The CAM3.0 is still imperfect and there are some spaces for further improvements. First, while we can control the tightness of clusters in the radius-fixed clustering, the computation time is much higher than K-means clustering. Because radius-fixed clustering forms clusters sequentially while K-means clustering determines clusters simultaneously, this is a need to integrate these two approaches and leverage their advantages. For example, we can use K-means clustering to initialize the radius-fixed clustering, though the uncertainty of K-means clustering remains as a concern. Second, as an alternative to pre-clustering, we may adopt robust linear programming (Ben-Tal and Nemirovski, 2000) to find convex hull with some randomness. Third, the identification of optimal simplex may be formulated as a Lasso-type vertex selection problem (Yuan and Lin, 2006), where the feature points with significant coefficients will be selected as true or highly probable vertices while determining the penalty and hyperparameter remain an unresolved problem. Fourth, except the clustering step, most other steps in the CAM3.0 pipeline are suitable for parallel computing.

To determine the effective scatter dimension, a critical dimension reduction step for reducing the subsequent computational complexity, we adopt deep-learning based strategies (Hu, et al., 2017; Yang, et al., 2019), where the eigenvalues of data covariance matrix are used to estimate possible number of sources via a fully-connected deep neural network. In the CAM3.0 pipeline, we use top 30 normalized singular values of data matrix as the inputs and extensive simulation datasets to train the deep neural networks. While this step aims to estimate the scatter dimension, we use twice of the estimated dimensions as the up limit for MDL based model selection.

To achieve convexity and shape preserved simplex visualization, based on the method introduced in debCAM, we further propose another projection strategy that can retain the distances among vertices. Specifically, first the order of vertices on the circle is estimated according to the distance among the vertices, and then the arc length is determined by the distance between the directly neighboring vertices (Seth and Eugster, 2016). The steps of the newly proposed projection are as follows:

1. For the *K* vertices of the (*K*-1)-simplex, find the three points (for example, *b, d*, and *f*) which form the triangle with the maximum area among all possible 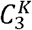 combinations exhaustively.
2. After finding out the three points, project them onto a circle temporarily (*b*′, *d*′, and *f*′), note that the arc length is not important in this step.
3. For the remaining *K*-3 points, sort them by the sum of distance to the three points (*b, d*, and *f*) detected in the first step decreasingly in a list.
4. According to the order after sorting in the third step, find the first point (for example, *a*), and compare the distance between this point (*a*) to the points not in the list (*b, d*, and *f*). Find out the two points which are the closest to the point (*d* and *f*), and then the point is projected onto the arc between the projection of the two points (*a*′ is between *d*′ and *f*′). Remove the point from the list.
5. Repeat the fourth step until there is no points left in the list.

Assign the arc length by the distance of the two end points of each arc before projection (the original space).

*Note: Supplementary information is available on the related website.*

## ACKNOWLEDGMENTS

This work was funded in part by the National Institutes of Health under Grants HL111362-05A1, HL133932, NS115658, and the Department of Defense under Grant W81XWH-18-1-0723 (BC171885P1).

## AUTHOR CONTRIBUTIONS

N/A.

