## Supplementary Information for "Determining molecular archetype composition and expression from bulk tissues with unsupervised deconvolution"

### Supplementary Tables

**Table S1** Counts of *a priori* markers enriched in CAM3.0-identified cell types (HBT)

| Brain span | S1  red | S2  blue | S3  cyan | S4  purple | S5  orange |
| --- | --- | --- | --- | --- | --- |
| Astrocyte (18*) | 15 | 0 | 0 | 0 | 0 |
| Mature oligodendrocyte (18*) | 0 | 10 | 0 | 0 | 0 |
| Neuron (13*) | 0 | 6 | 0 | 0 | 2 |
| Progenitor (25*) | 0 | 0 | 22 | 1 | 0 |

*count of probes measured in GSE25219 and linked to *a priori* marker genes

**Table S2** Counts of *a priori* markers enriched in CAM3.0-identified cell types (Braincloud)

| Braincloud | S1  red | S2  blue | S3  cyan | S4  green |
| --- | --- | --- | --- | --- |
| Astrocyte(33*) | 22 | 0 | 0 | 0 |
| Mature oligodendrocyte(23*) | 1 | 13 | 0 | 0 |
| Neuron(11*) | 0 | 3 | 1 | 3 |
| Progenitor(38*) | 3 | 0 | 15 | 0 |

*count of probes measured in GSE30272 and linked to *a priori* marker genes

**Table S3** Subtypes (counts of *a priori* markers) in brain tissues detected by (Xu, et al., 2013)

|  | Glia | Neuron |
| --- | --- | --- |
| FC(PC) | Astrocyte(18)  Mature oligodendrocyte(17) | Neuron(13) |
| CB | Astrocyte and Bergman glia(11)  Mature oligodendrocyte(9) | Inner golgi neuron(6)  Stellate/basket(5)  Granule(19)  Purkinje(9) |

**Table S4** Counts of *a priori* markers enriched in CAM3.0-identified cell types (PC)

|  | S1 | S2 | S3 |
| --- | --- | --- | --- |
| Astrocyte(18) | 13 | 0 | 0 |
| Mature oligodendrocyte(17) | 0 | 17 | 0 |
| Neuron (13) | 0 | 0 | 7 |

**Table S5** Counts *of a priori* markers enriched in CAM3.0-identified cell types (CB)

|  | S1 | S2 | S3 | S4 |
| --- | --- | --- | --- | --- |
| Astrocyte(11) | 5 | 0 | 0 | 0 |
| Mature oligodendrocyte(9) | 0 | 9 | 0 | 0 |
| Inner golgi (6) | 0 | 0 | 0 | 0 |
| Stellate/basket(5) | 0 | 0 | 0 | 1 |
| Granule(19) | 0 | 0 | 3 | 0 |
| Purkinje(9) | 0 | 0 | 0 | 8 |

**Table S6** Counts of marker proteins detected in LAD by csSAM and CAM3.0 and their overlaps

| Overlap | csSAM-NL  (83) | csSAM-FS  (102) | csSAM-FP  (115) |
| --- | --- | --- | --- |
| CAM3.0-NL1  (41) | 2 | 2 | 0 |
| CAM3.0-NL2  (50) | 30 | 1 | 0 |
| CAM3.0-FS  (46) | 0 | 26 | 0 |
| CAM3.0-FP  (42) | 0 | 1 | 26 |

**Table S7** Counts of marker proteins detected in AA by csSAM and CAM3.0 and their overlaps

| Overlap | csSAM-NL  (8) | csSAM-FS  (44) | csSAM-FP  (55) |
| --- | --- | --- | --- |
| CAM3.0-NL/FS  (283) | 3 | 16 | 0 |
| CAM3.0-FP  (50) | 0 | 0 | 46 |

**Table S8** Correlation coefficient between CAM3.0-estimation and pure measurement

| Correlation coefficient | FP Specimen 1 | FP Specimen 2 | FP Specimen 3 | FP Specimen 4 |
| --- | --- | --- | --- | --- |
| CAM3.0 in LAD | 0.8586666 | 0.6613852 | 0.8438751 | 0.8634448 |
| CAM3.0 in AA | 0.9193555 | 0.7554338 | 0.8634865 | 0.8820235 |

**Table S9** Correlation coefficients between estimated A matrix and ground truth A matrix of original CAM and improved CAM. The correlation coefficients of both versions are quite similar and high.

| Correlation coefficients | **Source 1** | **Source 2** | **Source 3** | **Average** |
| --- | --- | --- | --- | --- |
| **Original CAM** | 0.9797 | 0.9699 | 0.9845 | 0.9780 |
| **Improved CAM** | 0.9788 | 0.9690 | 0.9842 | 0.9773 |

**Table S10** Cosine similarities between estimated A matrix and ground truth A matrix of original CAM and improved CAM. The cosine similarities of both versions are quite similar and high.

| Correlation coefficients | **Source 1** | **Source 2** | **Source 3** | **Average** |
| --- | --- | --- | --- | --- |
| **Original CAM** | 0.9971 | 0.9908 | 0.9859 | 0.9913 |
| **Improved CAM** | 0.9972 | 0.9904 | 0.9885 | 0.9921 |

**Table S11** Period definition and sample number in Braincloud dataset. The definition of each period is from (Kang, et al., 2011). Though there some period of fetus with no samples, we still have enough samples for fetus at the other periods.

| Period | Description | Age (year) | Sample |
| --- | --- | --- | --- |
| 1 | Embryonic | -0.65 ~ -0.58 | 0 |
| 2 | Early fetal | -0.58 ~ -0.54 | 0 |
| 3 | Early fetal | -0.54 ~ -0.48 | 4 |
| 4 | Early mid-fetal | -0.48 ~ -0.42 | 9 |
| 5 | Early mid-fetal | -0.42 ~ -0.37 | 25 |
| 6 | Late mid-fetal | -0.37 ~ -0.27 | 0 |
| 7 | Late fetal | -0.27 ~ 0 | 0 |
| 8 | Neonatal and early infancy | 0 ~ 0.5 | 17 |
| 9 | Late infancy | 0.5 ~ 1 | 1 |
| 10 | Early childhood | 1 ~ 6 | 12 |
| 11 | Middle and late childhood | 6 ~ 12 | 4 |
| 12 | Adolescence | 12 ~ 20 | 49 |
| 13 | Young adulthood | 20 ~ 40 | 53 |
| 14 | Middle adulthood | 40 ~ 60 | 73 |
| 15 | Late adulthood | 60 ~ | 22 |

### Supplementary Figures

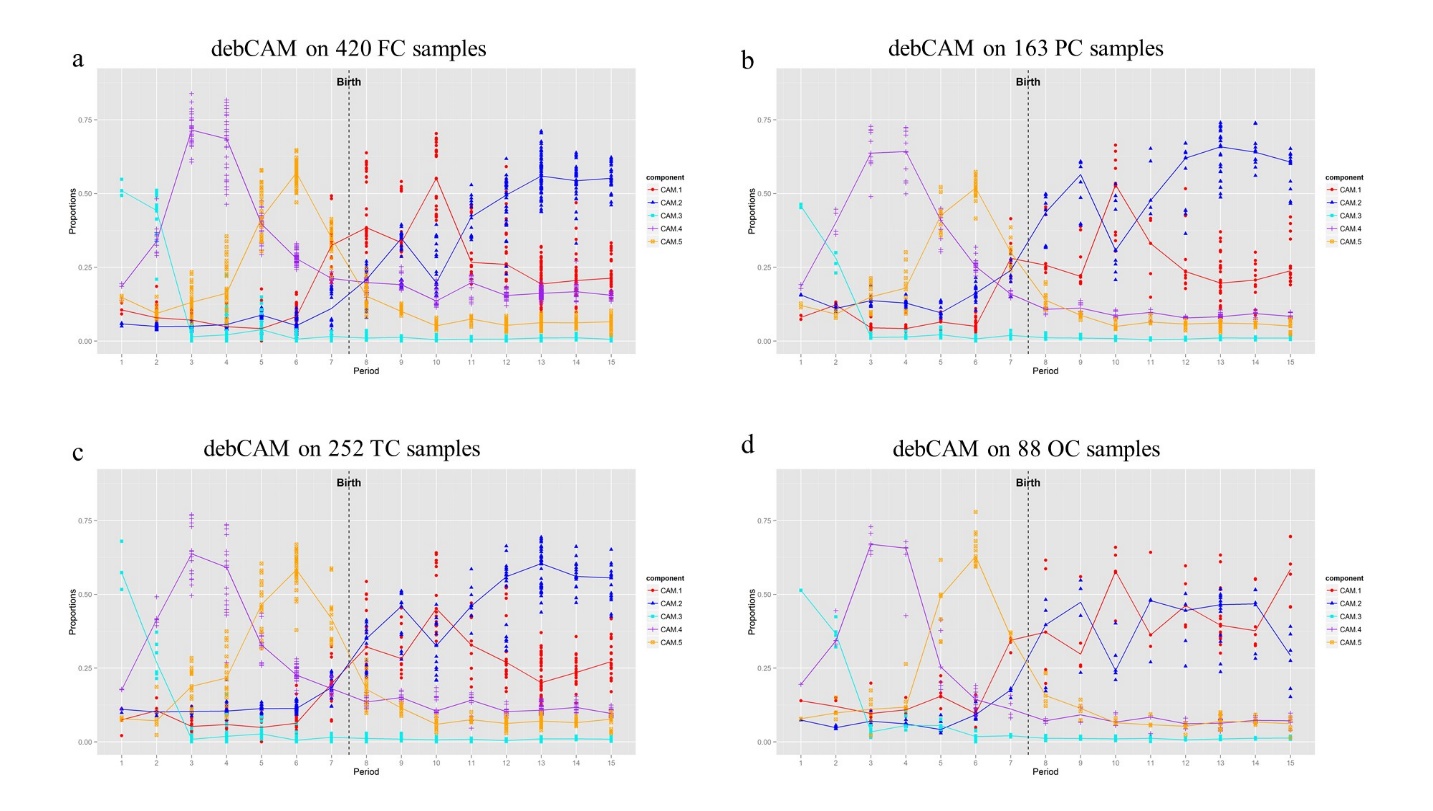

**Figure S1** Repopulation dynamics of distinctive subtypes/states during life span in each of four cortices estimated by CAM3.0 (implemented in the prototype software debCAM) applied to gene expression data from HBT cortex tissues.

| a PC  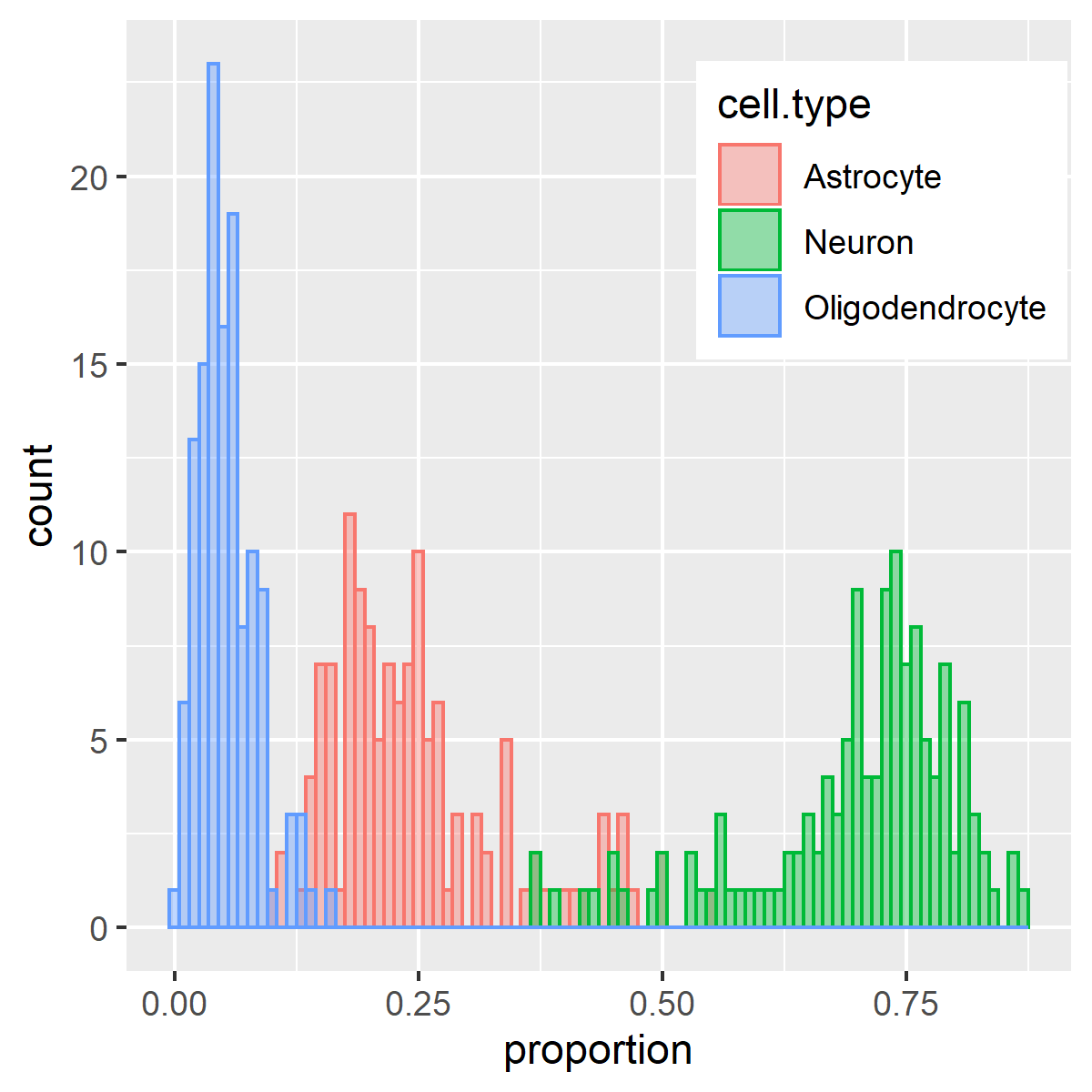 | b CB  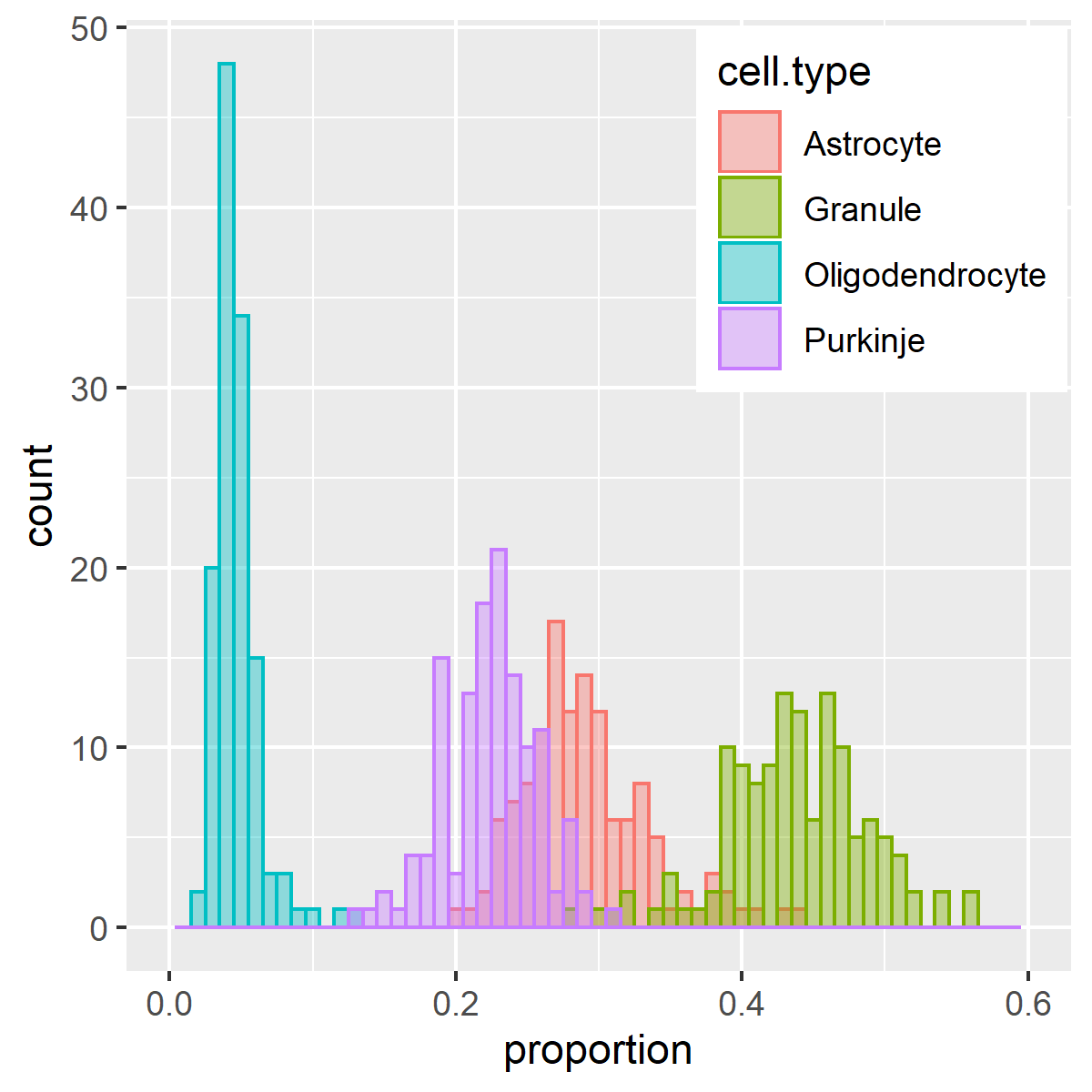 |
| --- | --- |
| **Figure S2** Histogram of estimated proportions of CAM3.0-identified cell types in parietal cortex and cerebellum (implemented in the prototype software debCAM). | |

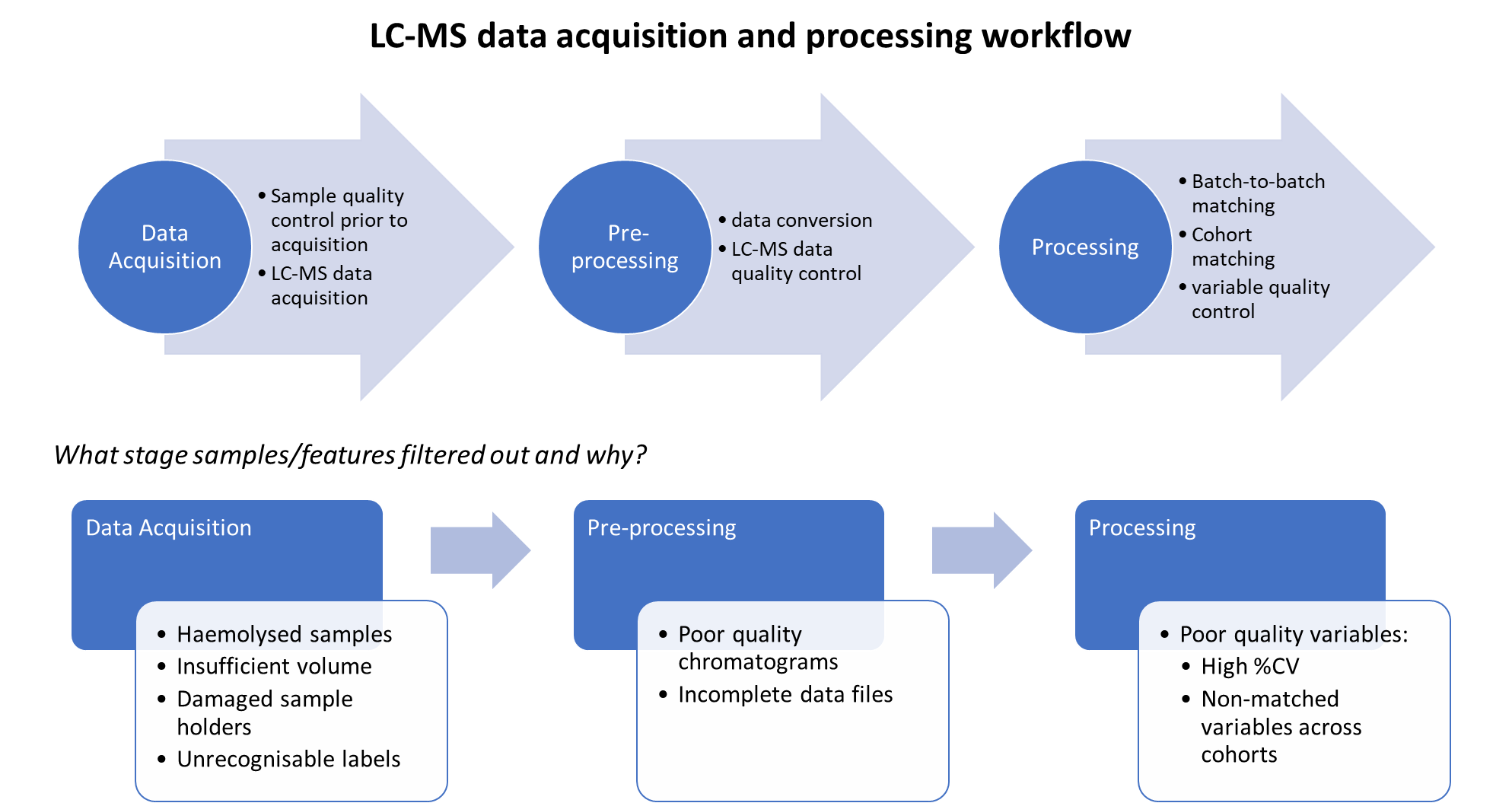

**Figure S3** Workflow of LC-MS data acquisition and preprocessing.

Kang, H.J.*, et al.* Spatio-temporal transcriptome of the human brain. *Nature* 2011;478(7370):483-489.

Xu, X., Nehorai, A. and Dougherty, J.D. Cell type-specific analysis of human brain transcriptome data to predict alterations in cellular composition. *Systems Biomedicine* 2013;1(3):0--1.
